# SIRT1-IDR-Driven Temporal Gating of Sequential Transcription-Factor Recruitment Coordinates Fasting Adaptation

**DOI:** 10.1101/2025.05.27.656293

**Authors:** Arushi Shukla, Sourankur Chakrabarti, Chinthapalli Balaji, Ullas Kolthur-Seetharam

## Abstract

Precise temporal control of gene expression is critical for metabolic adaptation, yet how transcriptional regulators coordinate dynamic responses to nutrient stress remains poorly understood. Here, we identify the intrinsically disordered region (IDR) of SIRT1, encoded by exon-2 (E2), as a key determinant of transcriptional timing during fasting. Using global and conditional SIRT1^ΔE2^ mouse model, which mimics an endogenous transcript variant, we demonstrate that loss of the N-terminal IDR disrupts the hierarchical engagement of transcription factors (TFs), leading to premature and exaggerated activation of gluconeogenic genes. Mechanistically, the IDR acts as a “conformational timer,” competitively gating SIRT1 interactions with CREB, FOXO1, and PPARα to ensure sequential TF recruitment and resolution. Absence of this regulatory module resulted in “runaway” CREB activity, aberrant hepatic glucose output, and impaired fasting-refeeding transitions, linking IDR-driven temporal control to systemic metabolic homeostasis. Our findings reveal a paradigm wherein IDRs in upstream metabolic sensors orchestrate transcriptional handovers, offering a molecular basis for the dynamic rewiring of gene networks during physiological transitions. This work expands the functional repertoire of disordered regions beyond TFs, positioning them as critical nodes in nutrient-responsive transcriptional circuits and potential targets for metabolic disorders.

## Introduction

Precise spatial and temporal regulation of transcription is essential for adapting gene expression during physiological transitions and environmental changes. In mammals, transcriptional complexity arises from cis-regulatory elements with multiple transcription factor (TF) binding sites and upstream co-regulators that modulate TF activity ^1–3^. While much is known about TF–DNA and TF– co-regulator interactions, emerging work highlights a critical role for intrinsically disordered regions (IDRs) within TFs that influence chromatin scanning, promoter selectivity, and target gene activation^4–6^.

IDRs have redefined the mechanistic underpinnings of cis-trans and trans-trans interactions, crucial for physico-chemical organisation of transcription, including spatial control via phase separation and condensates. Independently, while promoter specificity is dictated by DNA-binding domain (DBD), IDRs have been established as key determinants of promoter selectivity or promiscuity ^7–10^. Theoretical/Heuristic approaches, besides others, have clearly provided unique insights in this context. Importantly, thermodynamic frustration is known to contribute to transcriptional output by enabling TF interactions with multiple partners (Mediators, co-activators, co-repressors etc) to exert context-dependent regulation ^11–14^. However, most of our current understanding is limited to the role of IDRs in TFs with a paucity of information vis-à-vis regulation stemming from disorder-dependent dynamic interactions in upstream factors such as p300/CBP, SIRT1, and others. To reiterate, even though factors like KATs (TIP60, GCN5, p300, etc.) and KDACs (Sirtuins, HDAC1, HDAC3, etc.) are pivotal for transcription factor activity, synergistic or antagonistic interactions between them that are possibly mediated by IDRs in KATs/KDACs are less understood.

Despite this there are two key aspects which seem to have been largely unanswered in the field. Firstly, it is interesting to note that eukaryotic genomes have complex *cis*-regulatory landscapes (enhancers, promoters, insulators) and IDRs have been speculated to have co-evolved with these elements to manage increasing regulatory demands ^15–18^. But causal evidence to demonstrate this co-evolution in transcriptional machinery is non-trivial. Second and probably more important aspect is the bi-directional interplay between IDRs and physiology. This is key because most biological processes involve physiological transitions that entail transcriptional rewiring. Even though IDRs have been recognised as drivers of dynamic protein-protein interactions, necessary for transcription, if/how these influence or control temporal cascades of gene-expression remains poorly understood. Given this context, we hypothesize that IDRs in upstream regulatory factors act as conformational timers to not only orchestrate temporal gene expression but also to buffer aberrant cis-trans interactions on promoters with multiple TF binding sites.

Given this context, starvation offers an ideal model to investigate how intrinsically disordered regions (IDRs) in upstream regulators influence transcriptional timing in vivo. Physiological transitions such as development, immune responses, and metabolic adaptation require dynamic gene regulation by multiple transcription factors (TFs) acting in sequence or combination. Although overlapping regulatory inputs are well known, how TF modules are temporally ordered or exchanged remains unclear. The liver’s fasting response, particularly gluconeogenesis, involves TFs such as CREB, FOXO1, HNF4α, and the glucocorticoid receptor ^19–22^. While each is essential, their sequential activation during fasting is poorly understood. SIRT1, a NAD⁺-dependent deacetylase, plays a key role in this process. Unlike other metabolic sensors, SIRT1 contains extended N- and C-terminal IDRs. Its N-terminal IDR, encoded by exon 2, interacts with starvation-responsive TFs, but its in vivo role in coordinating transcriptional transitions is not fully known ^23–27^. Understanding this may reveal broader principles of dynamic transcriptional control during nutrient stress.

In this study, we examine the role of the SIRT1 N-terminal IDR, encoded by exon 2, in regulating transcriptional dynamics during fasting. Using a conditional SIRT1^ΔE2^ mouse model, we dissect its in vivo contribution to hepatic gene regulation and systemic glucose homeostasis. We find that the IDR temporally gates SIRT1 interactions with key starvation-responsive TFs, notably CREB. Loss of this region results in premature and amplified activation of gluconeogenic genes, increased pCREB occupancy on chromatin, and excessive hepatic glucose output, culminating in impaired fasting-refeeding transitions. These findings identify the SIRT1 IDR as a molecular timer that ensures transcriptional responses are appropriately phased. More broadly, this work highlights the regulatory potential of IDRs in upstream metabolic sensors, expanding their functional relevance beyond TFs to include fine-tuning of transcriptional networks during physiological transitions.

## Results

### Temporal regulation of gluconeogenesis and metabolic flux during starvation

The ability to mount an appropriate starvation response is critical not only for survival but also necessary to tune feed-fast cycles, which is inherently dependent upon metabolic plasticity, and is largely dictated by hepatic physiology. Among others, hepatic gluconeogenesis, is both essential for and diagnostic of anabolic-catabolic balance to sustain systemic metabolic needs. Despite decades of work that have identified upstream mechanisms that govern this, the temporal regulation of pyruvate utilisation and its channelling into gluconeogenesis remains insufficiently understood. Towards this, we assessed gluconeogenic potential, using PTT, as a function of duration of starvation in adult male mice, as indicated (Figure 1A and 1B). Our results revealed a significantly elevated glucose production at 24 hours of fasting when compared to 6 hours in response to the same amount of exogenously administered pyruvate. Although anticipated and consistent with current literature ^28–36^, these results offer direct experimental evidence to posit dynamic regulation of metabolite channelling during progressive fasting states, which is less understood.

**Figure 1:**
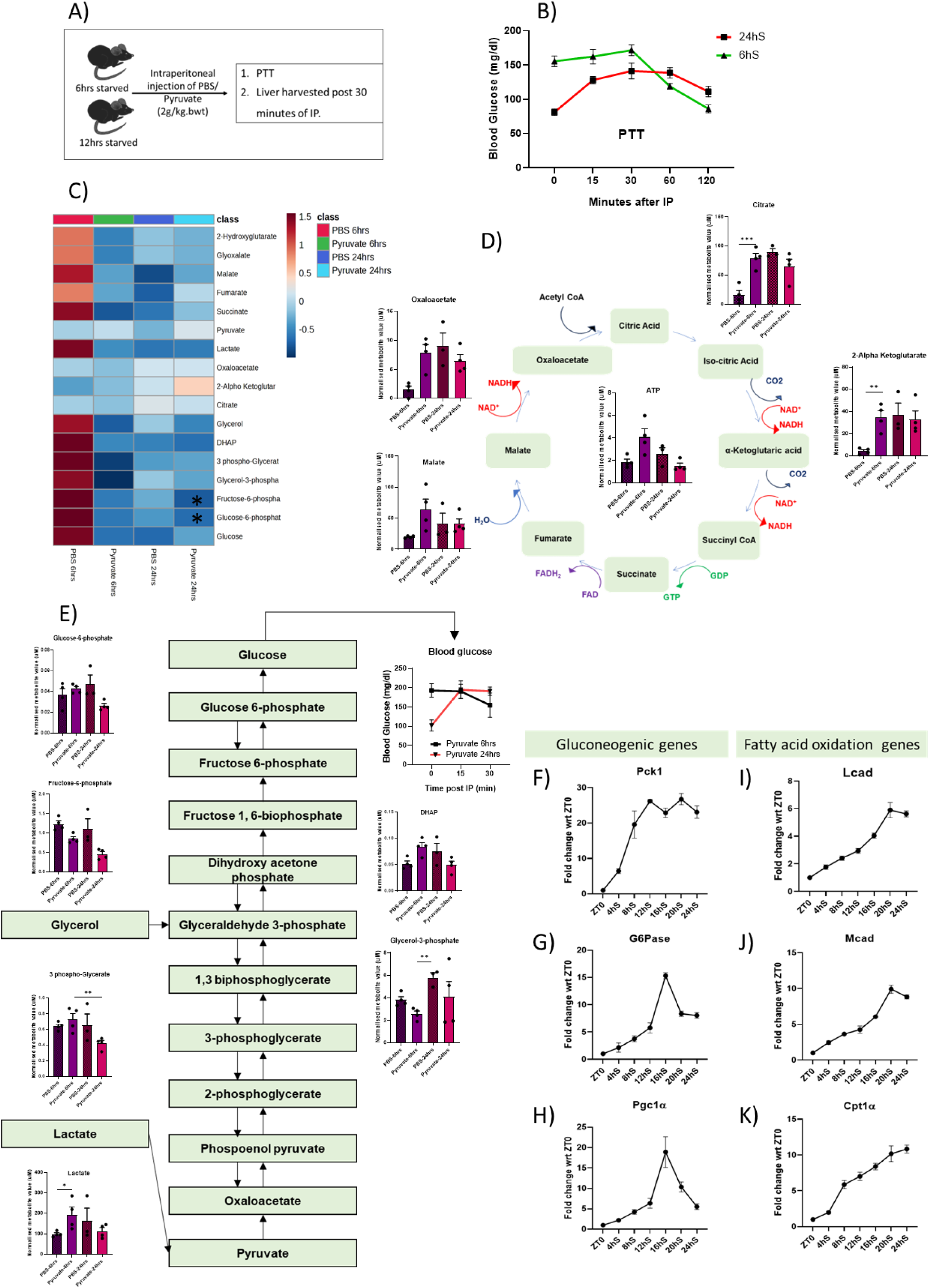
Gluconeogenesis and metabolic flux are temporally regulated during progressive starvation. **(A)** Schematic of the experimental design comparing 6-hour and 24-hour starvation conditions. **(B)** Pyruvate tolerance test (2 g/kg body weight) conducted at 6 h and 24 h of starvation; triangle markers (green line) represent 6 h, and square markers (red line) represent 24 h. The graph was plotted using GraphPad Prism 8.0. **(C)** Heatmap of normalized liver metabolite concentrations generated using MetaboAnalyst 5.0. **(D-E)** Schematic diagrams of the gluconeogenic pathway and TCA cycle annotated with normalized metabolite levels plotted on GraphPad Prism 8.0. **(F-K)** qPCR analysis of selected metabolic genes in livers from mice starved for indicated durations. Transcript levels were normalized to β-actin (N = 2, n = 3).

To gain further insights into differential metabolic channelling at 6- and 24-hours of starvation (6hS and 24hS), we analysed hepatic metabolome at baseline (PBS control) and in response to pyruvate administration (Figure 1C). As expected, the baseline metabolite profiles differed markedly between the 6hS and 24hS groups. Stable blood glucose and largely unchanged profiles of gluconeogenic intermediates suggested a predominant reliance on glycogenolysis at 6-hours of starvation. However, pyruvate bolus led to a significant increase in the levels of TCA cycle metabolites. In contrast, the 24hS group showed increased levels of tricarboxylic acid (TCA) cycle intermediates, indicating enhanced oxidative metabolism consistent with prolonged fasting, at baseline. It is interesting to note that 24-hour baseline profile was comparable to 6-hour pyruvate administration. Moreover, following pyruvate administration, there was a marked reduction in key intermediates such as fructose-6-phosphate, glucose-6-phosphate and Glycerol-3-phosphate, indicative of high pyruvate-dependent gluconeogenic potential (Figure 1 D and 1E). These striking, pyruvate-dependent changes are significant across pathways, especially for the glucose metabolism (Sup. Fig. 1A-1D). The differential re-wiring was further evident by changes in the ratios of key intermediates before and after pyruvate bolus (Sup. Fig. 1E-F). For example, ratios of many TCA metabolites to gluconeogenic pathway intermediates such as αKG/G6P and OAA/G6P were distinct at 6- and 24-hours of starvation. Importantly, this clearly illustrated biased utilization of substrates, which is dependent upon basal starvation-duration mediated molecular processes.

### Dynamic gene expression governs progressive starvation

Others and we have established the central role of transcriptional and post-transcriptional mechanisms in regulating starvation response during normal feed-fast cycles ^29,37–41^. However, the bi-directional interplay between hepatic metabolome and global gene expression cascades, and the cause-consequence mechanistic basis for precise temporal control of starvation response is much less understood. Towards this, we assessed for nodal genes whose expression is linked with glucose and fat metabolism across starvation phases. Gene expression analysis revealed a biphasic transcriptional response (Figure 1H), with an early phase (0–12 h) marked by induction of gluconeogenic genes, followed by a later phase (12–24 h) enriched for fatty acid oxidation (FAO) genes, and importantly correlated well with temporal change in hepatic metabolome. Further, we assayed for levels and/or post-translational modifications of transcription factors at these time points, as indicated. We specifically found that phosphorylated CREB (pCREB) peaked during the early starvation phase, while there was no change in the levels of FOXO1 and PPARα (Sup. Fig. 1H).

### Temporal control of SIRT1’s interactions by N-terminal disordered region

Emerging literature has demonstrated the importance of intrinsically disordered regions, especially in transcription factors, in mediating transient protein-protein interactions and their deterministic role of IDRs in dictating promoter selectivity ^5,7–9^. Further, it is necessary to note that there are no reports, which have shown relevance of IDR-dependent dynamic switch in transcription factor engagement in an in-vivo physiological setting, particularly during feed-fast cycles. Given this, we employed IUPred to compare presence of IDRs in proteins, which are central for mounting starvation response in the liver ^42^. We found that while CREB and FOXO1 were predicted to contain multiple disordered regions, nuclear receptors such as PPARα and RXR appeared largely structured (Figure 2A and Sup. Fig. 2A). However, it was interesting to note that co-regulators of transcription viz. CRTC2, PPARGC1A (PGC-1α) and SIRT1 displayed high degree and/or longer stretches of IDRs within them (Figure 2A and Sup. Fig. 2A).

**Figure 2:**
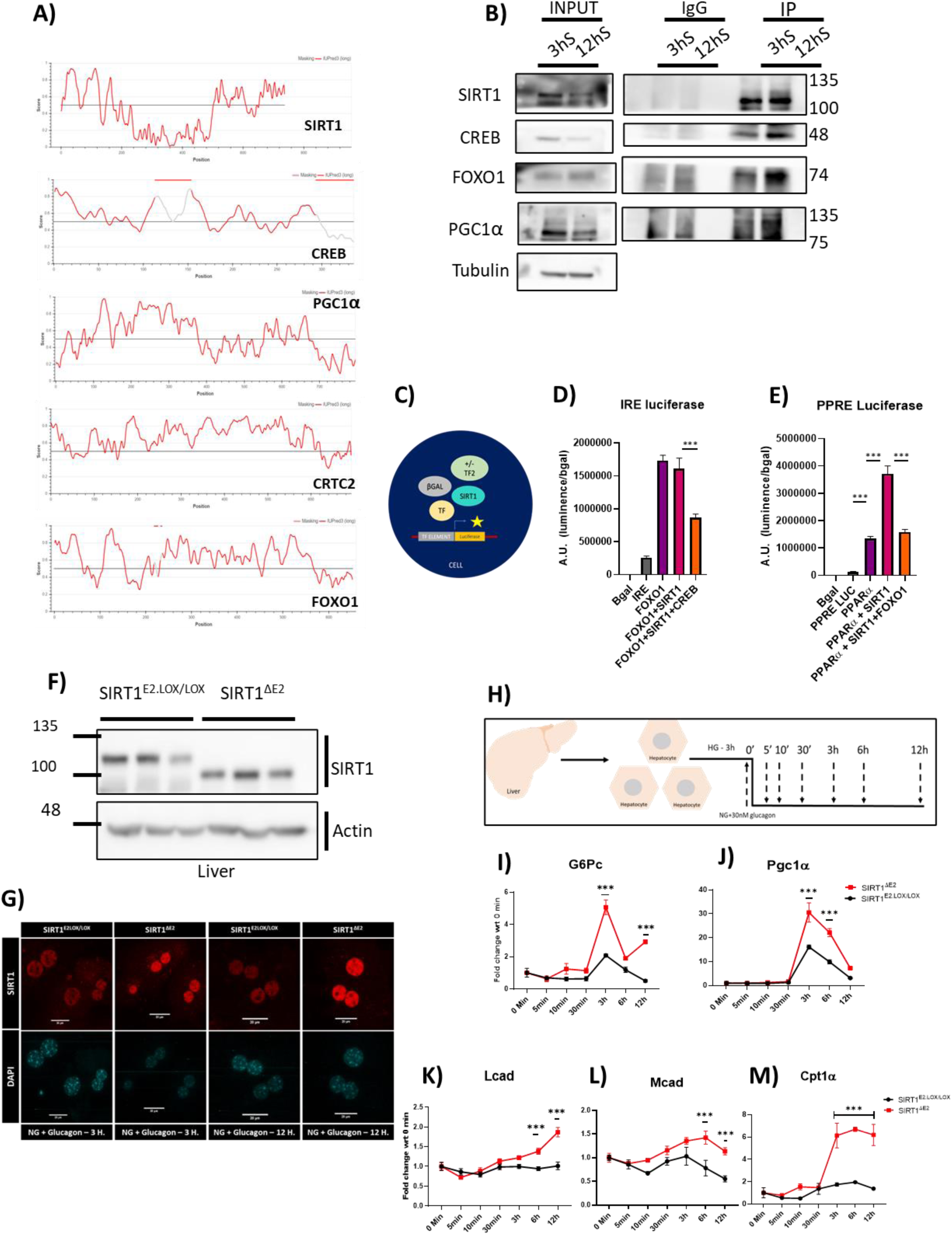
N-terminal intrinsically disordered region in SIRT1 regulates its interactions with TFs and regulates starvation transcriptional response. **(A)** Intrinsic disorder prediction profiles of indicated proteins generated using IUPred. The horizontal threshold line at 0.5 distinguishes disordered (above) from structured (below) regions. **(B)** Co-immunoprecipitation using SIRT1 antibody from liver lysates collected after 3 h and 12 h starvation, probed for CREB, FOXO1 and PGC1α. **(C)** Schematic of luciferase reporter assay design for assessing SIRT1-mediated transcriptional regulation. **(D-E)** Reporter assays to test competitive binding effects on SIRT1-mediated activation or repression of target promoters. **(F)** Western blots validation of SIRT1^ΔE2^ expression in the liver. **(G)** Representative immunofluorescence images confirming the nuclear localisation of SIRT1^ΔE2^ under starvation conditions. **(H)** Schematic of the primary hepatocyte culture and glucagon treatment timeline. Cells were harvested at indicated time points. **(I-M)** Time-course gene expression profiles of indicated glucagon-responsive genes in SIRT1^ΔE2^ and control hepatocytes. All graphs were plotted on GraphPad Prism 8.0, statistical analysis were done using 2-way anova; significance as *,p<0.05; **, p<0.005; ***, p<0.0005.

SIRT1 is known to act as a master regulator of transcription, during starvation, by modulating the activities of almost all the transcription factors and co-regulators ^43–49^. In this context, it was tempting to investigate if the interactions of SIRT1 with these nodal factors are dynamically remodelled in response to starvation, through co-immunoprecipitation from livers of mice starved for 3- and 12-hours (3hS and 12hS). As shown in Figure 2B and consistent with earlier literature, we indeed observed a starvation-duration-dependent rewiring of the SIRT1 interaction. Notably, there was a distinguishable change wherein the interactions with CREB, PGC1α and FOXO1 increased with prolonged fasting (Figure 2B). Given that the regulation of activity of key TFs converges at SIRT1, it led us to posit that competitive interactions may fine-tune SIRT1’s target selection and the transcriptional output downstream to these TFs.

To test this, we employed luciferase reporter assays using minimal TF-responsive elements (PPRE, IRE, CRE). In concordance with the literature, SIRT1 activated PPRE and IRE elements but suppressed CRE-driven transcription (Sup. Fig. 2B-2D). Moreover, TF competition assays showed that SIRT1-mediated activation of PPRE (PPARα-dependent) and IRE (FOXO-1-dependent) is antagonised by FOXO1 and CREB, respectively (Figure 2C-2E). This supported our hypothesis wherein SIRT1-TF selectivity drives competitive binding and hence preferential promoter-specific transcription. This is also consistent with our earlier report that illustrated the pertinence of the N-terminal region of SIRT1 encoded by exon-2 in mediating interactions, and is differentially spliced out in a tissue-specific manner.

### SIRT1 Exon 2 deletion enhances CREB-dependent transcription during starvation

Given that exon-2 encoded part is disordered and since IDRs have been shown to be involved in dynamic protein-protein interactions, we wanted to assess the importance of SIRT1-IDR in this context. Towards this, we generated a lox-E2-lox line to obtain conditional loss of exon-2 in SIRT1, as described in the methods section. We specifically wanted to investigate the importance of SIRT1-IDR-dependent engagement with TFs during dynamic physiological transitions, viz., starvation. As shown in Sup. Fig. 2F we floxed out exon-2 in young adults using tamoxifen-induced Cre activation and loss of exon-2 was confirmed across tissues (Sup. Fig. 2G-2H and Figure 2F).

Others and we have shown that although SIRT1 is predominantly nuclear, it shuttles between cytosol and the nucleus, which is also determined by glycosylation within the exon-2, particularly in a fed state^38^. Hence, we assessed the localisation of SIRT1^E2.lox/lox^ and SIRT1^ΔE2^ under starvation-mimicking conditions (no-glucose media supplemented with 30 nM glucagon) and found that both SIRT1^E2.lox/lox^ and SIRT1^ΔE2^ were predominantly localised in the nucleus (Figure 2G). Coupled with our earlier report wherein we had reported that lack of exon-2 did not affect catalytic activity, this prompted us to uncover if/how IDR mediated interactions determined fasting response at both molecular and physiological levels. It is important to note that transcriptional induction of key gluconeogenic genes (Pck1, G6pc1, and Pgc1α) was significantly higher in SIRT1^ΔE2^ hepatocytes and clearly indicated that loss of IDR in SIRT1 N-terminus modulates the magnitude of the transcriptional response by enhancing glucagon-induced gene activation (Figure 2H-M).

This further led us to investigate the temporal dynamics of CREB activation/functions and found rapid/heightened CREB phosphorylation at Ser133 in SIRT1^ΔE2^ hepatocytes in response to glucagon stimulation (Sup. Fig. 2I-J). Moreover, this was consistent with the in-vivo data from SIRT1^E2.lox/lox^ and SIRT1^ΔE2^ mice following 12-hour fasting (Sup. Fig. 2K). Together, these data indicate that the loss of SIRT1 exon 2 leads to precocious and amplified CREB activation.

Even though CREB is necessary to elicit starvation response, downstream to glucagon-PKA axis, mechanisms that govern temporal thresholding/restriction of CREB dependent transcription is poorly understood. Based on the above results, we assayed for SIRT1-CREB interactions since SIRT1 is known to inhibit CREB and its cofactor CRTC2. As shown in Figure 3A, co-immunoprecipitation assays revealed that SIRT1-bound CREB was dramatically lower in SIRT1^ΔE2^ compared to SIRT1^E2.lox/lox^ in primary hepatocytes under starvation conditions. Exon-2 dependent interaction and consequent effect on CREB activity was further substantiated by increased levels of pCREB in the unbound fraction from SIRT1^ΔE2^ hepatocytes (Figure 3B).

**Figure 3:**
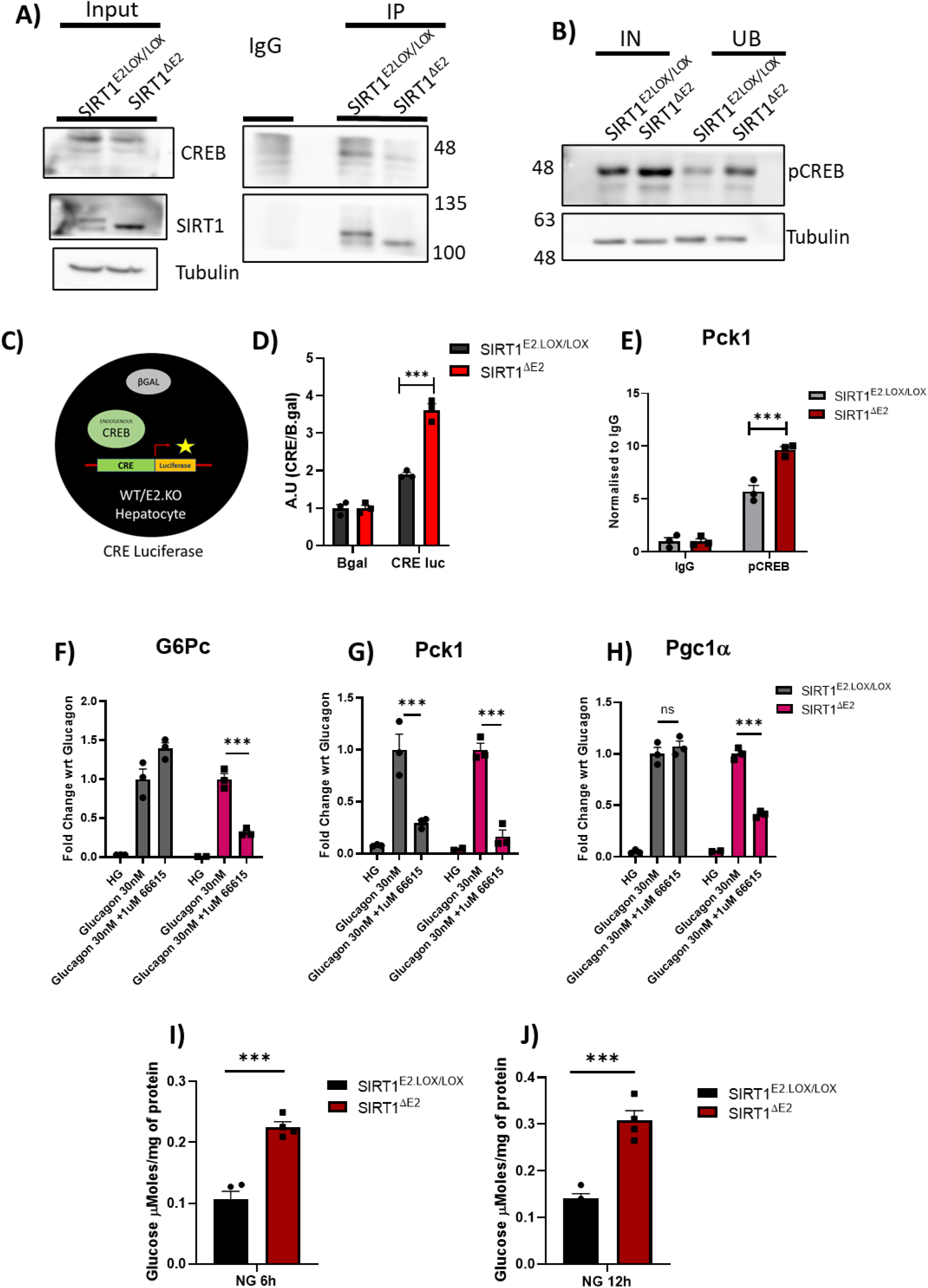
Exon 2 of SIRT1 is important for gatekeeping the starvation induced gluconeogenic response by regulating interaction between SIRT1 and CREB. **(A)** Co-immunoprecipitation (Co-IP) of SIRT1 followed by immunoblotting for SIRT1 and CREB in SIRT1^E2.LOX/LOX^ and SIRT1^ΔE2^ primary hepatocytes. **(B)** Input and unbound fractions from the Co-IP probed for pCREB. **(C)** Schematic of CREB-responsive element (CRE) luciferase reporter assay. **(D)** CRE reporter luciferase activity measured in SIRT1^E2.LOX/LOX^ AND SIRT1^ΔE2^ hepatocytes post glucagon stimulation. **(E)** ChIP-qPCR for pCREB occupancy on Pck1 promoter following glucagon treatment. **(F-H)** Relative expression of G6pc, Pck1, and Pgc1α after treatment with a CREB-specific inhibitor. **(I-J)** Gluconeogenesis assay was performed in SIRT1^E2LOX/LOX^ and SIRT1^ΔE2^ hepatocytes cultured under NG (6 h) and NG (12 h). All graphs were plotted on GraphPad Prism 8.0, one-way Anova was used for statistical testing (comparisons within the genotype only), significance as *,p<0.05; **, p<0.005; ***, p<0.0005.

Further this was substantiated by SIRT1 co-IP from the liver of progressively starved mice (3 h, 12 h, and 24 h). It was noted that across all time points, SIRT1^ΔE2^ mice exhibited consistently reduced CREB binding compared to SIRT1^E2.lox/lox^ (Sup. Fig. 3A–C), strengthening the finding that exon 2 is critical for maintaining SIRT1–CREB interaction during starvation.

### SIRT1 Intrinsically disordered region is important for gating CREB-Mediated transcriptional response

To determine the transcriptional consequences of this reduced interaction in a physiological setting, we performed CRE-luciferase reporter assays in SIRT1^E2.lox/lox^ and SIRT1^ΔE2^ hepatocytes following glucagon stimulation. CRE-Luciferase activity was significantly elevated in SIRT1^ΔE2^ cells, confirming heightened CREB-mediated transcription in the absence of exon-2 in SIRT1 (Figure 3C-D). More importantly, chromatin immunoprecipitation (ChIP) for pCREB showed increased occupancy at the Pck1 promoter and a similar trend of increase at G6pc in SIRT1^ΔE2^ hepatocytes (Figure 3E), demonstrating regulated recruitment of active pCREB, by SIRT1-exon-2, to gluconeogenic gene promoters.

In order to provide conclusive mechanistic evidence vis-à-vis SIRT1-IDR mediated modulation of pCREB dependent transcription, we pharmacologically inhibited CREB and PKA activity, as indicated. Pharmacological inhibition of PKA led to comparable reductions in gluconeogenic gene expression in both genotypes (Sup. Fig. 3E–G), indicating that enhanced CREB activity is not associated with altered upstream PKA signalling in SIRT1^ΔE2^. However, it was striking to find that treatment of hepatocytes with the CREB inhibitor, 666-15, led to a significantly blunted induction of Pck1, G6pc, and Pgc1α in SIRT1^ΔE2^ cells unlike in SIRT^E2.lox/lox^ hepatocytes (Figure 3F-H). This differential sensitivity to CREB inhibition clearly suggested that SIRT1^ΔE2^ hepatocytes exhibit heightened CREB activity akin to a “runaway” response.

The results described above prompted us to ascertain if this molecular rewiring led to a physiological output in terms of gluconeogenesis. As anticipated, SIRT1^ΔE2^ hepatocytes exhibited significantly elevated glucose production in response to both nutrient starvation and glucagon stimulation (Figure 3H-I and Sup. Fig. 3H-I). These together illustrated that loss of SIRT1-exon-2 promotes a hyperactive gluconeogenic state, even under mild starvation, besides providing confirmatory data on the regulatory control exerted by SIRT1-IDR on eliciting starvation response.

### SIRT1 IDR mediated gating is critical for systemic glycemic control and fed/fast cycles

Even though the results presented above clearly indicated the importance of SIRT1-IDR in programming gene expression during starvation, we wanted to investigate the in vivo physiological relevance of hepato-centric consequence of loss of exon-2 in SIRT1. As shown in Figure 4A, SIRT1^ΔE2^ mice exhibited increased body weight, compared to SIRT1^E2.lox/lox^, in the absence of any measurable difference in feeding behaviour (Figure 4A-B). Next, we assessed systemic glucose homeostasis through glucose tolerance test (GTT), as described in the methods section. SIRT1^ΔE2^ mice displayed markedly impaired glucose clearance, with significantly increased area under the curve (AUC), and a failure to return to basal glycaemic levels (Figure 4C-D). Further, we employed the pyruvate tolerance test (PTT) and found exaggerated hyperglycaemic response in SIRT1^ΔE2^ mice (Figure 4E). Dynamic monitoring of blood glucose levels during fasting also revealed consistently elevated glucose in SIRT1^ΔE2^ mice (Figure 4F), even in the absence of exogenous nutrient intake. Besides corroborating our in vitro and molecular data, these provide confirmatory evidence vis-à-vis the importance of SIRT1-IDR in orchestrating systemic metabolism.

**Figure 4:**
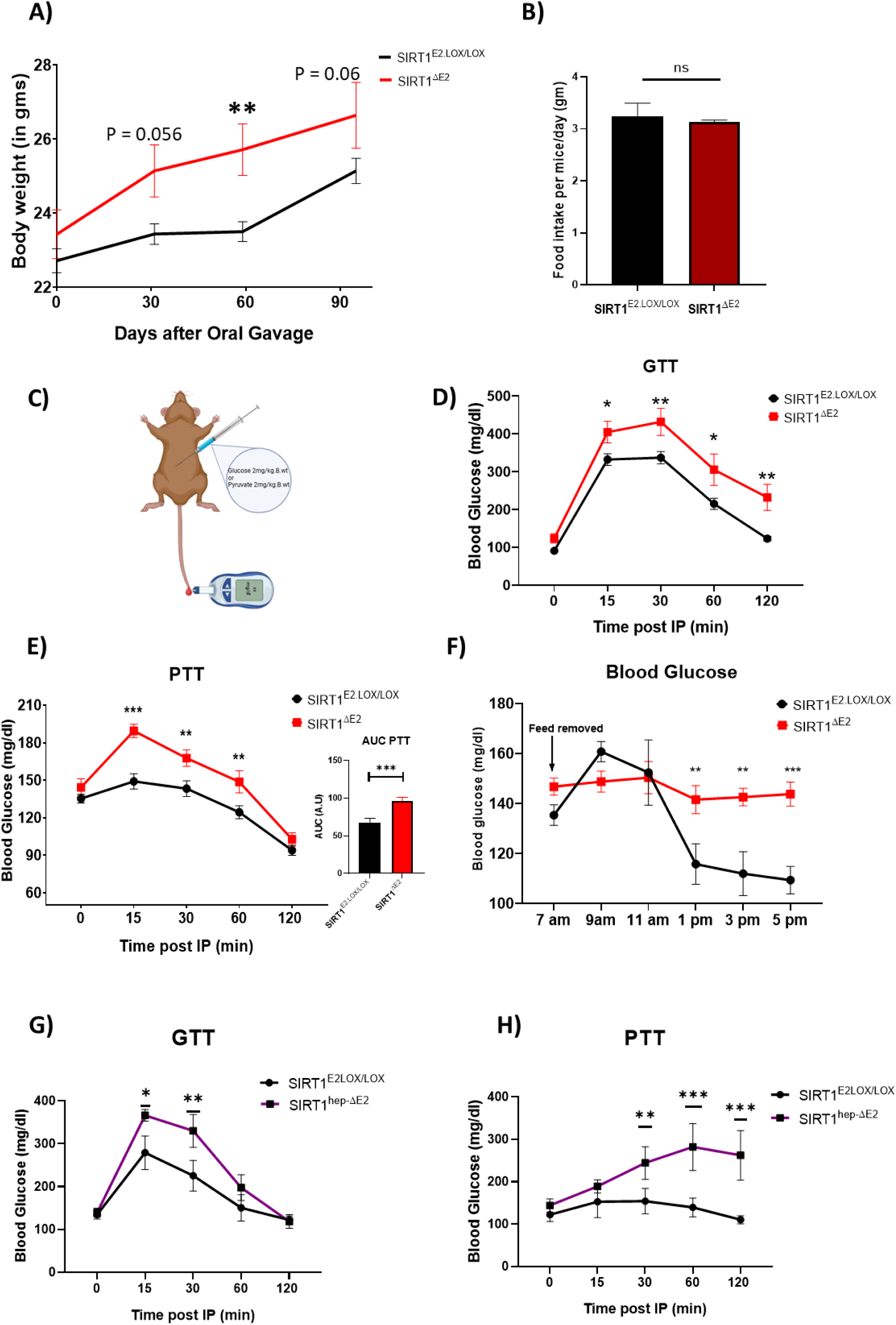
Loss of exon 2 in SIRT1 results in systemic glucose dyshomeostasis: **(A)** *Body weights of SIRT1^ΔE^*^2^ *and SIRT1^E^*^2^*^.LOX/LOX^ were measured under standard chow conditions. **(B)** Average daily food intake was quantified and compared between SIRT1^ΔE^*^2^ *and SIRT1^E^*^2^*^.LOX/LOX^.* **(C)** *Schematic illustrating the Intraperitoneal injection protocol for glucose and pyruvate tolerance tests. **(D-E)** GTT and PTT were performed on 10-14-week-old male mice; SIRT1^ΔE^*^2^ *and SIRT1^E^*^2^*^.LOX/LOX^ male mice following a 12-hour overnight fast. **(F)** Glucose measurements were taken every 2 hours from mice starting with food removal at ZT0. **(G-H)** GTT and PTT were conducted in 10-14-week-old male SIRT1^hep-ΔE^*^2^ *and SIRT1^E^*^2^*^.LOX/LOX^ mice, after a 12-hour overnight fast. N=3, n=3-5. All graphs were plotted in GraphPad Prism 8.0, Student’s t-test was used for statistical significance, *, p<0.05; **, p<0.005; ***, p<0.0005*.

Given the hyperactivation of the starvation-associated transcriptional program observed in SIRT1^ΔE2^ mice, we next examined their ability to transition into a fed-state response appropriately. To this end, mice were randomly assigned to either a 12-hour fasting group or a fasting group followed by 6 hours of refeeding. In wild-type animals, refeeding resulted in robust activation of AKT phosphorylation at Ser473 and degradation of full-length SIRT1, consistent with a normal fed-state transition. In marked contrast, SIRT1^ΔE2^ exhibited attenuated AKT phosphorylation and failed to degrade the SIRT1^ΔE2^ isoform (Sup. Fig. 4A-B), indicating impaired insulin signalling and an ineffective fed-state switch.

At the transcriptional level, SIRT1^ΔE2^ livers maintained elevated expression of fasting-induced genes, including *Pck1*, even following refeeding. Concurrently, the induction of canonical fed-responsive genes such as *Srebp1c* and *Fasn* was significantly blunted (Sup. Fig. 4C–H). Moreover, after prolonged starvation (24 hours) followed by short-term refeeding (3 hours), insulin receptor (InsR) transcription was significantly reduced in SIRT1^ΔE2^ mice (Sup. Fig. 4I), suggesting impaired insulin signalling. Together, these findings underscore the importance of the intrinsically disordered N-terminal region of SIRT1, demonstrating that it is indispensable for gating the transcriptional transition between fasting and feeding. Loss of this regulatory module leads to persistent activation of starvation-associated pathways, a failure to initiate fed-state transcriptional programs, and disruption of insulin signalling.

### Liver specific IDR loss disrupts glucose homeostasis / SIRT1 Exon2 deletion in liver drives metabolic imbalance

Others and we have earlier shown the importance of unravelling organ-system specific molecular mechanisms that drive organismal physiology, largely based on tissue specific knockouts or knockdowns. While IDRs have been shown to orchestrate transcription using in vitro cell culture models, there are no reports that have discovered the function of IDRs within specific tissues that impinge on whole body physiology. Hence, we generated liver-specific SIRT1 exon 2 knockout mice (SIRT1^hep-ΔE2^) by crossing SIRT1^E2.lox/lox^ animals with Albumin-Cre mouse line (Figure 4G-H). It was interesting to note that SIRT1^hep-ΔE2^ animals phenocopied the SIRT1^ΔE2^ mice vis-à-vis GTT and PTT. These exciting results indicated that functions of SIRT1-IDR in the liver contributes to the systemic metabolic dysregulation and illustrates tissue-autonomous modulation of SIRT1-TF interactions, mediated by the non-catalytic domain encoded by exon-2, is necessary for starvation response at an organismal level. These results reveal, for the first time, that a non-catalytic IDR within SIRT1 mediates liver-specific transcriptional programs that are essential for maintaining whole-body metabolic balance. By demonstrating that loss of the SIRT1-IDR disrupts the temporal coordination of transcription factor interactions during starvation, our findings uncover a broader principle: that intrinsically disordered regions within metabolic sensors enable dynamic transcriptional handover in response to physiological cues. This highlights a novel layer of regulatory control through which disordered protein domains shape tissue-specific gene expression to sustain systemic homeostasis.

## Discussion

Temporal engagement of transcription factors, which is critically dependent upon both cis-binding of upstream regulatory elements and trans interactions, is essential for orchestrating programmed reversible physiological transitions e.g., during fed-fast-refed cycles ^2,16,50,51^. Even though endocrine and metabolic signalling cascades that impinge on TF activity are well established, foundational principles of protein-protein interactions that mediate kinetic gating of promoter-selective recruitment of TFs during dynamic metabolic toggling remain poorly understood ^28,31,52–56^. Our findings reveal that loss of IDR within SIRT1 disrupts metabolic fidelity, leading to premature transcriptional responses, aberrant glucose homeostasis, and impaired fasting-refeeding transitions. By bridging the gap between IDR dependent interaction dynamics and physiological adaptation, we posit that SIRT1 IDR acts as a critical arbiter of transcriptional timing, besides providing hitherto unknown insights into molecular origins of metabolic disorders rooted in dysregulated interactions.

The biphasic transcriptional response during starvation—early gluconeogenic gene induction followed by fatty acid oxidation—aligns with the dynamic rewiring of SIRT1’s interactome. The IDR-mediated competitive binding of TFs (e.g., CREB vs. PPARα) provides a mechanistic basis for promoter selectivity and temporal resolution of gene expression. This is consistent with emerging paradigms that IDRs enable multivalent, context-dependent interactions. While considerable attention has been given to the roles of IDRs in transcription factors ^6,15,57^, comparatively fewer studies have explored their function within transcriptional coactivators or metabolic regulators. Recent findings in p300/CBP suggest that IDRs in coactivators support multivalent interactions and enhance chromatin engagement ^58–61^, underscoring the relevance of such mechanisms in transcriptional regulation.

Our work extends this paradigm to metabolic sensors, demonstrating that SIRT1’s N-terminal IDR governs transcriptional handovers during nutrient stress. This aligns with reports showing that IDRs in nutrient-responsive enzymes, such as ULK, RAPTOR, CAMKII etc, enable allosteric regulation, PTM-based regulation and substrate selection ^62–69^. The observation that SIRT1^ΔE2^ hepatocytes exhibit “runaway” CREB activity parallels findings in p53, where disordered regions prevent promiscuous DNA binding by enforcing transient TF interactions. This dysregulation led to elevated expression of gluconeogenic genes and systemic glucose dyshomeostasis, establishing a mechanistic link between temporal interaction specificity and metabolic output. Such kinetic control ensures transcriptional fidelity, preventing aberrant activation under fluctuating nutrient conditions.

Notably, liver-specific deletion of SIRT1’s exon 2 recapitulated systemic glucose dysregulation, highlighting the tissue-autonomous role of this IDR in maintaining whole-body homeostasis. The failure of SIRT1^ΔE2^ mice to transition effectively from fasting to feeding states, marked by blunted insulin signalling and persistent gluconeogenic activation, suggests that the IDR acts as a conformational timer to buffer aberrant transcriptional cascades. This aligns with broader evidence that IDRs in nutrient sensors like SIRT1 co-evolved with regulatory networks to integrate metabolic cues and transcriptional outputs.

The liver’s ability to toggle between fed and fasted states hinges on transcriptional and metabolic plasticity, a process tightly regulated by nutrient-sensing machinery ^30,41,70–73^. Our findings reveal that SIRT1’s IDR is indispensable for this dynamic adaptation. During fasting, SIRT1-IDR ensures precise temporal engagement with CREB to activate gluconeogenesis, while its competitive interactions with PPARα and FOXO1 prevent premature lipid oxidation. Conversely, refeeding demands rapid suppression of fasting programs and reactivation of anabolic pathways, such as insulin-mediated glycogenesis. Previously we had shown the importance of SIRT1 in activating the insulin-dependent fed response; however ^38^, failure to resolve fasting signals in SIRT1^ΔE2^ mice mirrors metabolic inflexibility and insulin resistance, underscoring the IDR’s role as a rheostat for transcriptional transitions. Our work thus positions SIRT1’s IDR as a critical node in the transcriptional circuitry that gates hepatic metabolic identity, ensuring coherent shifts between energy storage and mobilisation. Notably, this study significantly advances our understanding of transcriptional regulation under nutrient stress by identifying the N-terminal IDR of SIRT1 as a critical regulatory element that acts as a molecular gatekeeper, modulating SIRT1’s interactions with key TFs like CREB, FOXO1, and PGC-1α. These are consistent with emerging literature that propose IDRs act as conformational switches.

Taken together, our study bridges a critical gap in understanding how disordered regions in upstream regulators (e.g., deacetylases) govern TF activity beyond their catalytic functions, offering a mechanistic blueprint for how disordered domains sustain metabolic resilience across fed-fasted-refed cycles. SIRT1’s IDR ensures fidelity in transcriptional handovers during starvation; we propose a model where IDRs serve as kinetic checkpoints to prevent runaway responses. Future work could explore whether this mechanism extends to other physiological transitions, such as immune activation or circadian rhythms, where dynamic TF engagement is paramount.

## Materials and methods

### Animal Experiments

All the experiments and protocols were done in accordance with the Institutional Animal Ethics Committee (IAEC) guidelines and approvals. Mice were housed at standard housing conditions of 12h light/dark cycles (at TIFR Mumbai, CCMB, IISER Pune animal houses). The temperature was maintained at 20° C ± 2 and humidity at 50 ± 20. They were fed normal chow diet and had free access to autoclaved drinking water, unless otherwise mentioned. Mice strains include C57BL/6NCrl, SIRT1^E2.LOX/LOX^ (generated as described), ERT2^CRE/CRE^ (Jax Stock ID : 008463), Alb^CRE/CRE^ (Jax Stock ID : 018961), SIRT1^E2.LOX/LOX^; ERT2^CRE/CRE^, SIRT1^E2LOX/LOX^; Alb^CRE/CRE^. All experimental animals were euthanised by cervical dislocation and the relevant tissues were isolated and snap frozen in liquid nitrogen, unless stated otherwise. SIRT1-Lox-E2-Lox mouse was generated by Turku Center for Disease Modeling University of Turku, Turku, Finland as a paid service.

### Tamoxifen administration for generating conditional whole body SIRT1 EXON 2 knockout model (SIRT1^ΔE2^)

To obtain whole-body conditional knockout of Exon 2 of SIRT1, 2-month-old male mice which were homozygous for E2 LOX and ERT2 Cre with the genotype SIRT1^E2.LOX/LOX^; ERT2^CRE/CRE^, were administered Tamoxifen by oral gavage. Tamoxifen was reconstituted in corn oil at a concentration of 20mg/ml by dissolving at 37 °C, and 4 doses of 100mg/kg body weight were administered on alternate days. Knockout was confirmed from tail clip DNA after 7 days from the last dose. Animals were used for experiments as per the paradigms used and explained below and in relevant sections.

### Generation of SIRT1 Exon 2 liver knockout

To obtain Liver-specific knockout of exon 2 of SIRT1, the SIRT1^E2.LOX/LOX^ were crossed with albumin Cre mice where the CRE is under the albumin promoter and hence the KO is restricted to the liver. The mice line was maintained as heterozygous SIRT1 ^E2LOX/+^; ALB ^cre/+^ and with the help of tail clip genotyping the SIRT1^hep-ΔE2^ (LKO) and SIRT1^E2.LOX/LOX^ (WT) animals were selected for the experiments described later.

### Glucose tolerance test

For glucose tolerance test (GTT), mice were starved for 12 hours and body weight were measured, baseline blood glucose was measured from tail prick using a glucometer (AccuCheck active) labelled as zero minutes. 2g/ kg body weight of glucose was injected intraperitoneally (I.P). Subsequently, the blood glucose was monitored at 15 mins, 30 mins, 60 mins, and 120 mins, post the glucose I.P injection. The data was plotted using GraphPad-Prism 8.0.

### Pyruvate Tolerance Test

For the pyruvate tolerance test (PTT), mice were starved for 6,12 or 24 hours, depending on the experiment and body weight was measured, baseline blood glucose was measured from tail prick using a glucometer (AccuCheck active) labelled as zero minutes. 2g/ kg body weight of pyruvate was injected intraperitoneally (I.P). Subsequently, the blood glucose was monitored at 15 mins, 30 mins, 60 mins, and 120 mins post the pyruvate I.P injection. The data was plotted using GraphPad-Prism 8.0.

### Primary hepatocyte culture

Male SIRT1^ΔE2^ and SIRT1^E2LOX/LOX^ mice (12–16 weeks old) were anaesthetised by intraperitoneal injection of Thiopentone (Neon Laboratories Ltd., Mumbai, India). Once sedation was confirmed, the mice were disinfected with 70% ethanol. Liver perfusion was performed through the inferior vena cava/portal vein using 30 mL of Hank’s Balanced Salt Solution (HBSS) (5.33 mM potassium chloride, 0.44 mM KH2PO4, 4.16 mM NaHCO3, 137.93 mM NaCl, and 0.338 mM Na2HPO4, pH adjusted to 7.4). The solution was supplemented with 5.5 mM glucose (Sigma-Aldrich G8769), 25 mM HEPES (USB 16926), and 100 mM EGTA (Sigma-Aldrich E3889). Following perfusion, the liver was digested using a digestion medium containing DMEM-Low Glucose (Sigma-Aldrich D5523), 15 mM HEPES, antibiotic-antimycotic (Anti-Anti), and collagenase type IV (Sigma-Aldrich C5138). The digested liver tissue was carefully minced and filtered through a 70 µm cell strainer to obtain a hepatocyte suspension. The cell suspension was then centrifuged at rcf value of 50 g for 5 minutes at 4 °C. The resulting pellet was washed thrice with high-glucose DMEM (25 mM glucose; Sigma-Aldrich D7777) supplemented with 10% fetal bovine serum (Gibco 16000044). The cells were counted using a haemocytometer and plated onto collagen-coated (Sigma-Aldrich C3867) 100 mm/ 6 well culture dishes at a density of 2X 10^6^ or 3 x 10^5^ cells per plate, respectively. The cells were incubated at 37 °C in a humidified atmosphere with 5% CO2 to allow for adherence. Six hours post-plating, the medium was replaced to remove non-adherent cells and replaced with high-glucose DMEM without any serum addition.

### Glucagon treatments in hepatocytes

Primary hepatocytes were isolated as described above, and the cell medium was changed to DMEM high glucose without serum for 12 hours. To expose hepatocytes to starvation-like conditions, the medium was changed to DMEM-No glucose media (supplemented with glutamine2mM and pyruvate 2mM) with the final concentration of glucagon at 30nM for the time points as mentioned. Post-treatment, the cells were collected in RIPA buffer or TRIzol as per the experiment’s requirement. The hepatocytes were metabolically synchronised for pulsatile glucagon experiments by replenishing the medium with HG-DMEM three hours before media change, followed by media change into DMEM-NO Glucose (± glucagon). Every media change was preceded by two phosphate buffer saline washes (PBS) (Thermo, #88512) to remove the residual media.

### CREB inhibitor (666-15) and PKA inhibitor (H89) treatments for hepatocytes

Primary hepatocytes were isolated as described above, and the cell medium was changed to DMEM high glucose without serum for 12 hours. To assess the effect of CREB and PKA inhibitor the hepatocytes were metabolically synchronised by replenishing the medium with HG-DMEM three hours before media change, followed by media change into the following: HG, DMEM-NO Glucose + glucagon (30nM), DMEM-NO Glucose + glucagon (30nM) + 666-15 (1µM) and DMEM-NO Glucose + glucagon (30nM) + H89 (1µM and 10µM). Cells were collected after 3 hours of treatment. Every media change was preceded by two phosphate buffer saline washes (PBS) (Thermo, #88512) to remove the residual media. Post-treatment, the cells were collected in RIPA buffer or TRIzol as per the experiment’s requirement.

### Plasmid transfection and luciferase assay in hepatocytes

Primary hepatocytes were isolated from Male SIRT1^ΔE2^ and SIRT1^E2LOX/LOX^ mice and plated as described above. Plasmid transfections were done using the JetPrime transfection reagent (Polyplus, #101000027) for 24-48 hours as per the manufacturer’s protocol. After 6 hours of plating, hepatocytes were transfected with a plasmid featuring the CREB promoter upstream of the luciferase gene, a kind gift from Prof. Mark Montminy, along with a β-galactosidase expressing plasmid. After 36 hours of transfection, the cells were treated with DMEM-NG + 30nM glucagon for 3 hours. The hepatocytes were washed with chilled PBS and harvested in passive lysis buffer (Promega # E194A) for 15 mins, followed by centrifugation at 12000 rpm. The supernatant was collected for the BCA estimation, β-Gal assay and the luciferase assay. β-Gal assay was performed using the ONPG (Sigma #N1127) method, where 30µL of supernatant is incubated with 50µL of ONPG (4mg/ml) at 37 °C for 20-30 mins. 100µL of stop solution (1M Na2CO3) was added to wells, and the absorbance was measured at 420nm. For the luciferase assay, 30µL of supernatant was taken in a black plate to this 100µL of luciferase assay system reagent (Promega #E1501) was added. The counts for the luminescence were measured in the TriStar LB941 (Berthold technologies) and normalised to the β-Gal activity and represented as fold change.

### Mass spec Sample Preparation

#### Polar Non-Derivatized Metabolite extraction: -

A known weight of liver tissue was homogenised in 75% ethanol containing internal standards: C13-Glucose (10 µM; Cambridge Isotopes, #CLM-1396) and L-Tryptophan (1.5 µM; Cambridge Isotopes, #DLM-615862). The samples were vortexed for 3 minutes, incubated at 80 °C for 3 minutes, and subsequently transferred to ice for 5 minutes. Following incubation on ice, the samples were centrifuged at 17,000 g for 10 minutes at 4 °C.

An equal volume of the resulting supernatant from all samples was transferred to fresh Eppendorf tubes and dried using a speed vacuum concentrator. The dried samples were stored at -20 °C until mass spectrometry analysis was performed.

#### Polar Derivatized (TCA) Metabolites extraction

A measured quantity of liver tissue was homogenized in 75% ethanol containing internal standards: **D4-**succinic acid (3 µM; Cambridge Isotopes #DLM-2307). The mixture was vortexed for 3 minutes, incubated at 80 °C for 3 minutes, and then incubated on ice for 5 minutes. After cooling, the samples were centrifuged at 17,000 g for 10 minutes at 4 °C. The supernatant was collected in equal volumes into fresh Eppendorf tubes and dried using a speed vacuum concentrator.

The dried samples were re-suspended in a 1:2 ratio of H₂O and 1 M N(3-dimethylaminopropyl)-N’-ethylcarbodiimide (prepared in 13.5 mM pyridine buffer, pH 5.0). The samples were mixed by tapping, followed by the addition of 150 µL of 0.5 M O-benzyl hydroxylamine (prepared in 13.5 mM pyridine buffer, pH 5.0). The samples were incubated at 25°C for 1 hour.

After incubation, 350 µL of ethyl acetate was added, and the mixture was shaken for 10 minutes. The samples were centrifuged at 3,000 g for 5 minutes at 4 °C, and the supernatant was transferred to a fresh tube. This ethyl acetate extraction process was repeated two more times. The collected supernatant was then dried using a speed vacuum concentrator. The final dried samples were stored at -20 °C until analysis by mass spectrometry.

#### Mass spectrometry Analysis: -

Dried samples were analysed using an Agilent 6546 Quadrupole Time-of-Flight (QTOF) mass spectrometer operating in Auto-MS/MS acquisition mode, coupled with an Agilent 1290 Infinity II UHPLC system. Liquid chromatography (LC) separation was performed using a Synergi 4 μm Fusion-RP 80 Å LC column (100 × 4.6 mm, Ea; catalogue no: 00D-4424-E) with a Gemini guard column (Phenomenex, 4 mm × 3 mm; catalogue no: KJ0-4282).

For polar non-derivatized metabolites, chromatographic separation was conducted at a flow rate of 0.5 mL/min. The mobile phase composition for positive ion mode consisted of solvent A: 99.9% (v/v) H₂O + 0.1% formic acid, and solvent B: 99.9% (v/v) methanol (MeOH) + 0.1% (v/v) formic acid. For negative ion mode, solvent A was 100% (v/v) H₂O with 5 mM ammonium acetate, while solvent B was 100% (v/v) acetonitrile (ACN). The gradient elution program was as follows: 0–3 min, 5% solvent B; 3–10 min, 5–60% solvent B; 10–11 min, 60–95% solvent B; 11–14 min, 95% solvent B (wash step); 14–15 min, 95–5% solvent B; and 15–21 min, 5% solvent B (re-equilibration step).

For polar derivatized metabolites, data acquisition was performed in positive ion mode with a flow rate of 0.5 mL/min. The mobile phase consisted of solvent A: 99.9% (v/v) H₂O + 0.1% formic acid, and solvent B: 99.9% (v/v) MeOH + 0.1% (v/v) formic acid. The LC run time was 30 min, with the following gradient elution program: 0–3 min, 0% solvent B; 3–10 min, 0–5% solvent B; 10–11 min, 5–60% solvent B; 11–18 min, 60–95% solvent B; 18–19 min, 95% solvent B (wash step); 19–22 min, 95–60% solvent B (re-equilibration step); 22–25 min, 60–5% solvent B; and 25–30 min, 5–0% solvent B.

The autosampler and column temperatures were maintained at 10 °C and 40 °C, respectively. Mass spectrometric data were acquired using a dual Agilent Jet Stream electrospray ionization (AJS ESI) source in both positive and negative ion modes. The MS operating parameters were as follows: drying gas temperature, 320 °C; sheath gas temperature, 320 °C; drying gas flow, 11 L/min; sheath gas flow, 11 L/min; nebulizer pressure, 45 psi; capillary voltage, 3 kV; nozzle voltage, 1 kV; and fragmentor voltage, 150 V. Precursor ion selection was performed with a maximum of 10 precursor ions per cycle, and collision energy was set at 5 eV.

### Data Analysis and Quantitation

LC-MS data processing was performed using Agilent MassHunter Qualitative Analysis 10.0. Identified peaks were manually verified based on retention times and fragment ions obtained from MS/MS spectral analysis. A mass accuracy threshold of 10 ppm was applied to all analytes across all experiments described in this study. Metabolite quantitation was conducted by calculating the area under the curve (AUC) of each analyte relative to its corresponding internal standard. The resulting values were subsequently normalised to the total protein content within liver tissue samples to ensure accurate absolute concentration measurements.

### RNA isolation and quantitative PCR (qPCR)

Total RNA was isolated from tissue or cells using TRIzol following the manufacturer’s instructions. Briefly, TRIzol was added to cells or tissues and homogenised using a pestle. Following 5 minutes of incubation at room temperature, 0.2 ml of chloroform was added per 1 ml of TRIzol. After shaking, samples were incubated for 2 mins and centrifuged at 12000 rpm for 15 mins at 4 °C. The aqueous phase was collected and transferred into a new microcentrifuge tube, and isopropanol was added in the ratio of 1 to 2 volumes of TRIzol, followed by 3-4 hours of incubation at -20 °C. Samples were centrifuged at 12000 rpm for 15 minutes and RNA pellets were washed using 70% ethanol. After washing, the RNA pellets were dried at 37 °C for 3-5 minutes and resuspended in the desired volume of DEPC water. The concentration and purity of RNA samples were measured using UV spectroscopy (by measuring the absorbance of samples at 260 and 280 nm).

1-5 μg RNA was utilised for cDNA preparation using Random Hexamer and SuperScript™ IV Reverse Transcriptase following the manufacturer’s protocol. Quantitative PCR was performed using the LightCycler® 96 (Roche) and the KAPA SYBR® FAST qPCR Master Mix (2X) Kit following the manufacturer’s guidelines. The expression of target genes was normalised with actin transcripts levels.

### Lysate preparation

Cells or tissue were homogenised in ice-cold RIPA buffer (50mM Tris pH 8.0, 150mM NaCl, 0.1% SDS, 0.5% Sodium deoxycholate, 1% Triton X-100, 0.1% SDS) supplemented with 1 mM PMSF, 1.5XProtease inhibitor cocktail, 1.5XPhostop and incubated for 10-20 minutes with brief vortexing, occasionally. The lysates were sonicated using Bioruptor® Pico (Diagenode) (5 cycles for cells and 10 cycles for tissues with 30 s ON / 30 s OFF at 4ᵒC) followed by centrifugation at 13000 rpm for 15 minutes at 4 °C. Supernatants were transferred into fresh microcentrifuge tubes and used for estimating protein concentration using BCA (Bicinchoninic Acid) Protein estimation assay as per the manufacturer’s instruction. Samples were boiled at 95 °C for 10 mins after adding Laemmli sample buffer (50 mM Tris-Cl pH 6.8, 2% SDS, 0.1% bromophenol blue, 10% glycerol, 100 mM DTT).

### Western Blotting

40-60 μg of equal protein were run on Sodium dodecyl-sulfate polyacrylamide gel electrophoresis (SDS-PAGE) and transferred to polyvinylidene fluoride (PVDF) membrane. After blocking with 5% Milk in Tris-buffered saline with 0.1% Tween (TBST) for 90 minutes, desired proteins were probed by overnight incubation of the membrane with primary antibodies (diluted in 5% BSA in TBST) at 4 °C with continuous rocking. Following three washes with TBST at room temperature, the membranes were incubated with horseradish peroxidase-conjugated secondary antibodies for one and a half hours at room temperature. Membranes were washed thrice with TBST at room temperature, and the bands of proteins were visualised in the GE Amersham Imager 600 instrument (General Electric) using a chemiluminescent-based detection kit. The band intensities of targeted proteins were measured using ImageJ and normalised with that of the loading control. Expression of actin or tubulin, or Histone H3 were utilised as a loading control, as mentioned in the respective figures.

### Immunoprecipitation (IP)

50ug of liver tissue from the mice as per the experiment or plated hepatocytes as per the treatment were collected in PBS, washed twice and lysed in TNN buffer (50mM Tris-cl pH 7.5, 150mM Nacl, 0.9% NP40), along with 1.5X PIC, 1.5X Phosstop and 1.5mM PMSF and incubated for 15 minutes. The lysate was centrifuged at 12000rpm at 4 °C. Supernatant was carefully separated and protein A beads were used for preclearing at 4 °C at slow rotor (5 rpm) for 1.5 hours. Following preclearing, input was separated and 3-4mg of protein was used for IP. Control antibody (Rabbit IgG) and SIRT1 were used for IP. Immunoprecipitation was allowed for 8-10 hours at 4 °C on at slow rotor followed by pull down with Prot A magnetic beads. After 3-4 hours, the unbound fraction was collected and the beads were proceeded for washing. Post the washes, the beads were magnetised, all the residual buffer removed, and laemmli buffer was added to the beads, vortexed for 5 seconds and boiled for 15 minutes. After boiling, the beads were magnetised again, and the supernatant was carefully separated. All the fractions were analysed as per the experiments need and mentioned in the relevant results section.

### Chromatin Immunoprecipitation (ChIP) from hepatocytes

#### Crosslinking

Hepatocytes (2×10^6^) were plated and in 100mm collagen-coated cell culture-treated Petri dishes. On the second day post-plating, both SIRT1^E2.Lox/Lox^ and SIRT1^ΔE2^ cells were treated with starvation mimicking media (NG+glucagon 30nm). After 3 hours, of treatment, cells were washed twice with ice-cold PBS and crosslinked with 1% formaldehyde for 8 minutes at room temperature. Crosslinking was quenched with 125 mM glycine for 5 minutes, followed by three washes with chilled PBS. Cells were scraped in cold PBS, collected by centrifugation at 1000 rpm for 5 minutes, the supernatant was discarded, and the resulting pellet was stored at −80 °C for further processing.

#### Lysis and sonication

Frozen cell pellets were thawed and resuspended in 600 µL of ice-cold Cell Lysis Buffer (CLB; see Sup. Table 1). After incubation on ice for 10 minutes, samples were centrifuged at 3500 rpm for 5 minutes at 4 °C, and the supernatant was discarded. The pellet was resuspended in 500 µL of CLB, vortexed briefly, and incubated on ice for another 5 minutes. This step was repeated once more with 300 µL of CLB. The final pellet was resuspended in Nuclear Lysis Buffer (NLB), incubated on ice for 10 minutes, and then diluted with 500 µL of ChIP Dilution Buffer. Samples were snap-frozen in liquid nitrogen and thawed on ice prior to sonication.

Sonication parameters were optimized to obtain chromatin fragments suitable for transcription factor (TF) ChIP. Post-sonication, samples were centrifuged at 13,000 rpm for 15 minutes at 4 °C, and the supernatant (sheared chromatin) was collected for immunoprecipitation and quality assessment.

### Reverse Crosslinking (for Chromatin Assessment)

To assess chromatin quality, 20 µL of the sheared chromatin was diluted with 180 µL of ChIP Dilution Buffer. To this, 8 µL of 5 M NaCl was added, and the mixture was incubated at 65 °C for 12–20 hours. RNase A (20 µL of 10 mg/mL) was added, vortexed, and incubated at 37 °C for 2 hours, followed by the addition of 5 µL of Proteinase K and incubation at 45°C for 2–4 hours. DNA was purified using a phenol: chloroform extraction (1:1), followed by a chloroform wash and ethanol precipitation with 2 volumes of chilled ethanol and 25 µL of 3 M sodium acetate. Precipitated DNA was centrifuged at 13,000 rpm at 4 °C for 30 minutes, washed with 70% ethanol, and air-dried. DNA was finally resuspended in 20 µL of nuclease-free water and assessed for concentration, purity, and fragment size before proceeding with ChIP.

### Bead Preparation

Protein A magnetic beads were washed with PBST (PBS + 0.1% Tween-20), then resuspended in ChIP Dilution Buffer supplemented with 10 µL of 10 mg/mL tRNA and 1% BSA. Beads were blocked by incubation at 4 °C with gentle rotation for 4 hours.

### Chromatin Immuno-precipitation (ChIP)

Approximately 35–50 µg of chromatin per immunoprecipitation (IP) was used. Blocked beads were added for pre-clearing at 4 °C for 2 hours with gentle rotation. After pre-clearing, 5% of the input was reserved, diluted to 200 µL with ChIP Dilution Buffer, and stored at −20 °C. The remaining pre-cleared chromatin was split into two aliquots: one incubated with rabbit IgG (negative control) and the other with anti-pCREB antibody. Antibody binding was performed overnight (10–12 hours) at 4 °C with gentle rotation. Subsequently, blocked beads were added to each tube and incubated for an additional 4 hours.

Beads were sequentially washed twice with the following buffers, each for 5 minutes at 4°C: Low Salt Buffer, High Salt Buffer, Lithium Chloride Buffer, and finally Tris-EDTA Buffer. Following washes, 100 µL of Elution Buffer was added, and the beads were vortexed and incubated at 65°C for 30 minutes. The supernatant was collected, and the elution was repeated with an additional 100 µL of Elution Buffer. Supernatants were pooled, and 8 µL of 5 M NaCl was added to both IP and input samples for reverse crosslinking, followed by overnight incubation at 65 °C.

### DNA Purification and qPCR

Reverse crosslinking was followed by RNase A and Proteinase K digestion, as described previously. Glycogen blue and 25 µL of 3 M sodium acetate were added, and DNA was precipitated overnight at −20 °C. After ethanol precipitation and drying, the DNA pellet was resuspended in 50 µL of nuclease-free water. The immunoprecipitated DNA was analysed by qPCR using primers for target gene loci.

### Gluconeogenesis assay

Hepatocytes (50,000 cells/well) isolated from SIRT1^E2.Lox/Lox^ and SIRT1^ΔE2^ mice were seeded in collagen-coated 24-well plates. Following adherence, cells were treated according to the experimental conditions—either with normal glucose (NG) medium alone or NG supplemented with 30 nM glucagon—for 6 or 12 hours. Prior to treatment, cells were thoroughly washed with PBS to remove residual serum or media components.

At the end of the treatment period, supernatants were collected for glucose measurement, and the corresponding wells were used to scrape cells for protein quantification by BCA. The glucose concentration in the culture supernatant reflects the extent of hepatocyte-mediated gluconeogenesis. Glucose levels were quantified using the Glucose Colourimetric Detection Kit (EIAGLUC, Invitrogen) following the manufacturer’s protocol. This assay is based on the enzymatic oxidation of glucose by glucose-oxidase, producing hydrogen peroxide, which in the presence of horseradish peroxidase (HRP) reacts with a chromogenic substrate to yield a coloured product measurable at 560 nm.

For quantification, assay standards provided in the kit were serially diluted to prepare the following concentrations: 32 mg/mL, 16 mg/mL, 8 mg/mL, 4 mg/mL, and 2 mg/mL. In each well, 20 µL of either sample or standard was added in duplicate. To this, 25 µL of HRP solution was added, followed by 25 µL of chromogenic substrate, and finally 25 µL of 1× glucose oxidase solution. The reaction mixtures were incubated at room temperature for 30 minutes.

Absorbance was measured at 560 nm using a microplate reader. A standard curve was generated from the known glucose standards, and sample concentrations were calculated accordingly.

### Immunofluorescence in Hepatocytes

Hepatocytes (50,000 cells/well) isolated from SIRT1^E2.Lox/Lox^ and SIRT1^ΔE2^ mice were seeded in collagen-coated coverslips. Following adherence, cells were treated according to the experimental conditions—either with High glucose (HG-25mM glucose) medium or no glucose (NG) supplemented with 30 nM glucagon—for 3 hours. After the treatment, cells were washed with chilled PBS and were then immediately fixed in 4% formaldehyde (methanol-free) (Thermo Scientific Cat. No # 28908) for 20 min at room temperature. Post fixation cells were washed gently with 1X PBS thrice and blocked for 1 hour in 5% BSA in TBST (0.05% Triton). Cells were then incubated with SIRT1 antibody (CST) at a concentration of 1:1000 overnight. Cells were washed thrice with 1X PBS and incubated with Alexa Fluor-conjugated secondary antibodies (Thermo-Fisher Scientific, AF 488 or AF 647) for 1 hour at room temperature. Cells were washed three times with 1X PBS and stained with DAPI (1:1000 of 1mg/ml). The coverslips were mounted in Vectashield on a glass slide and kept in place using transparent nail paint. They were taken for imaging post-drying in the FV3000 confocal system at 60X magnification with oil (Olympus).

## Quantification and statistical analysis

Data are expressed as means ± standard error of means (SEM). Statistical analyses were done using Graph Pad Prism (version 8.0). Student’s t-test and ANOVA were used to determine statistical significance. A value of p ≤ 0.05 was considered statistically significant. *p ≤ 0.05; **p ≤ 0.05; ***p ≤ 0.005.

## Acknowledgements

We thank Dr. Mark Montminy, for sharing the CRE reporter luciferase construct with us. We are grateful to TIFR Mumbai, IISER Pune and CCMB animal house for housing and maintaining our animals. We particularly thank Dr. Shital Suryavanshi, Dr. Kalidas N. Kohale, Dr. M Jerald Mahesh Kumar, Chetan Sable, S Prashanth and all the animal house staff for their invaluable help with animal handling and animal experiments. We extend our acknowledgement to UK lab members for their critical inputs and discussion during the study.

## Author contribution

**Conceptualization: A.S., U.K-S., Methodology: A.S., S.C., C.B., U.K-S., Investigation: A.S., S.C., Visualization: A.S., S.C., C.B., Supervision: U.K-S., Writing—original draft: A.S., U.K-S.**

## Data and Material availability

All data needed to evaluate the conclusion of this paper are present in the paper and/or the supplementary material.

## Supplementary Information

**Supplementary Figure 1.**
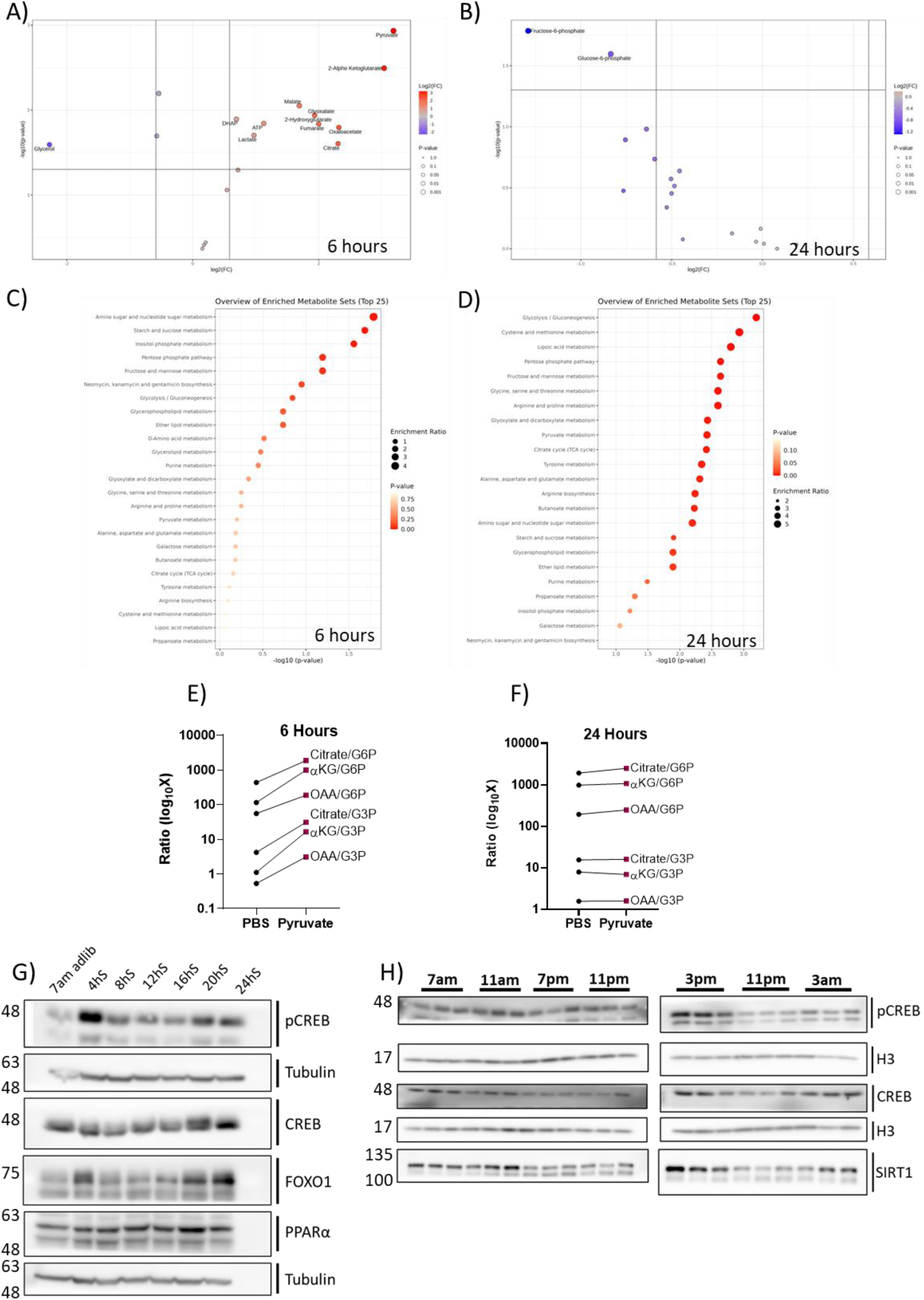
(Sup. Fig. 1): Metabolic signatures of temporal starvation in mouse liver. (A-B) Volcano plots depicting differentially abundant metabolites after PBS vs. pyruvate administration at 6 h and 24 h, respectively. **(C-D)** Pathway enrichment analysis for the respective groups based on metabolomics data using MetaboAnalyst 5.0. **(E-F)** Logarithmic increase in the ratio of metabolites is depicted between PBS and Pyruvate conditions at 6 h and 24 h, respectively. **(G)** Representative western blots showing levels of pCREB, CREB, FOXO1, PPARα, SIRT1, tubulin, and H3 at the starvation time points. **(H)** Protein expression analysis from the livers of mice subjected to time-restricted feeding (TRF) for 15 days and sacrificed at indicated zeitgeber times (ZT).

**Supplementary Figure 2.**
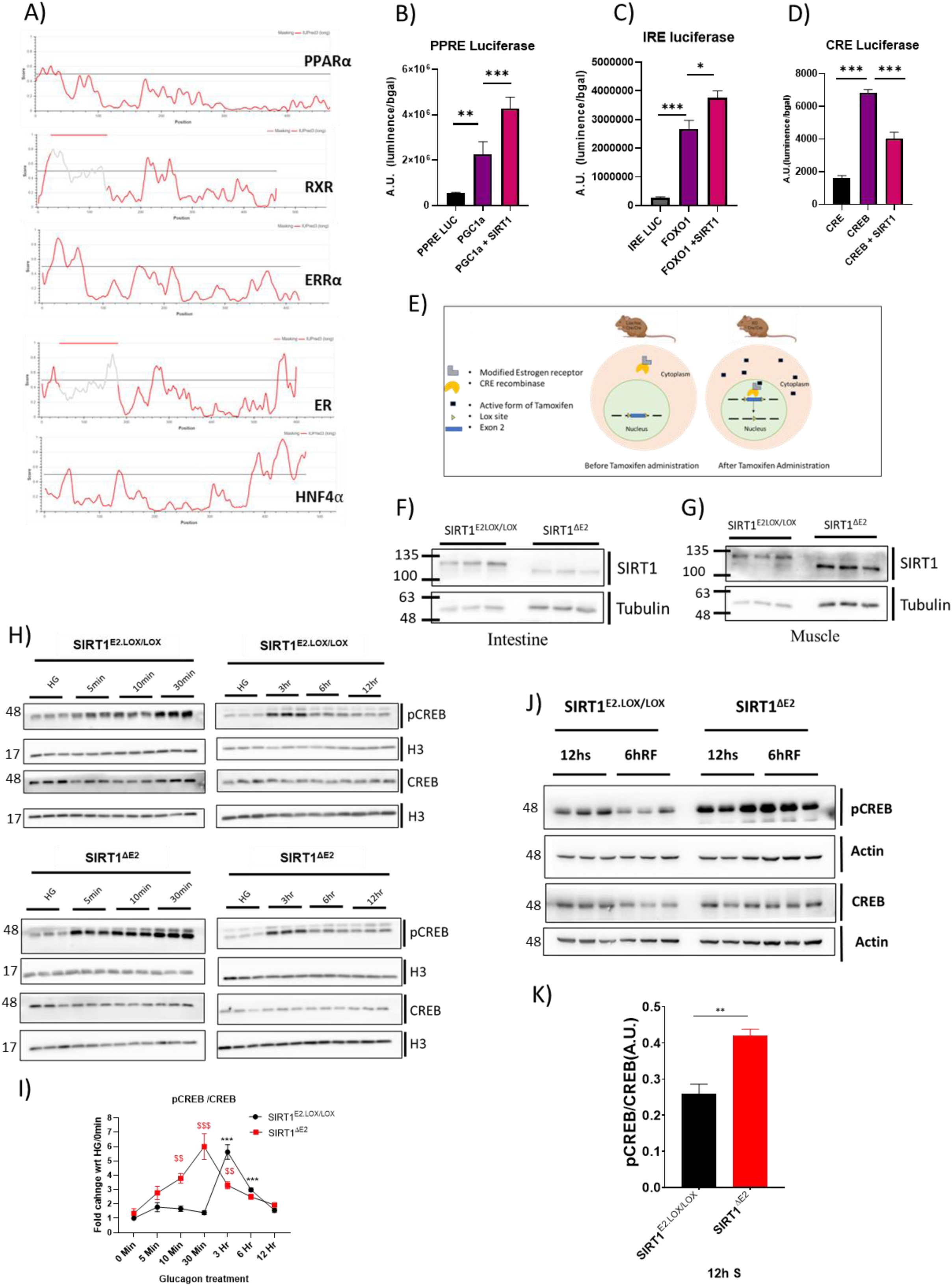
(Sup. Fig. 2): Molecular and Functional Validation of SIRT1^ΔE2^ and Its Impact on CREB Signalling. **(A)** Intrinsic disorder prediction profiles of indicated proteins generated using IUPred. The horizontal threshold line at 0.5 distinguishes disordered (above) from structured (below) regions. **(B-D)** Effect of SIRT1 overexpression on activity of CRE, IRE and PPRE reporter constructs. **(E)** Schematic showing the mechanism of tamoxifen-induced Cre recombination and generation of the SIRT1^ΔE2^ variant. **(F-G)** Western blots validation of SIRT^ΔE2^ expression in the intestine and muscle tissues respectively. **(H)** Western blot showing levels of total CREB, phosphorylated CREB (pCREB), and histone H3 in hepatocytes treated with glucagon (30 nM) for the indicated time points under nutrient-deprived (NG) or control (HG) conditions. **(I)** Quantification of pCREB levels in SIRT1^E2.LOX/LOX^ and SIRT1^ΔE2^, hepatocytes, analysed using one-way ANOVA relative to HG controls**. (J)** Western blot of liver lysates from mice after 12 h starvation and 6 h refeeding to assess CREB and pCREB levels in vivo. **(K)** Quantification of pCREB levels normalized to loading control.

**Supplementary Figure 3.**
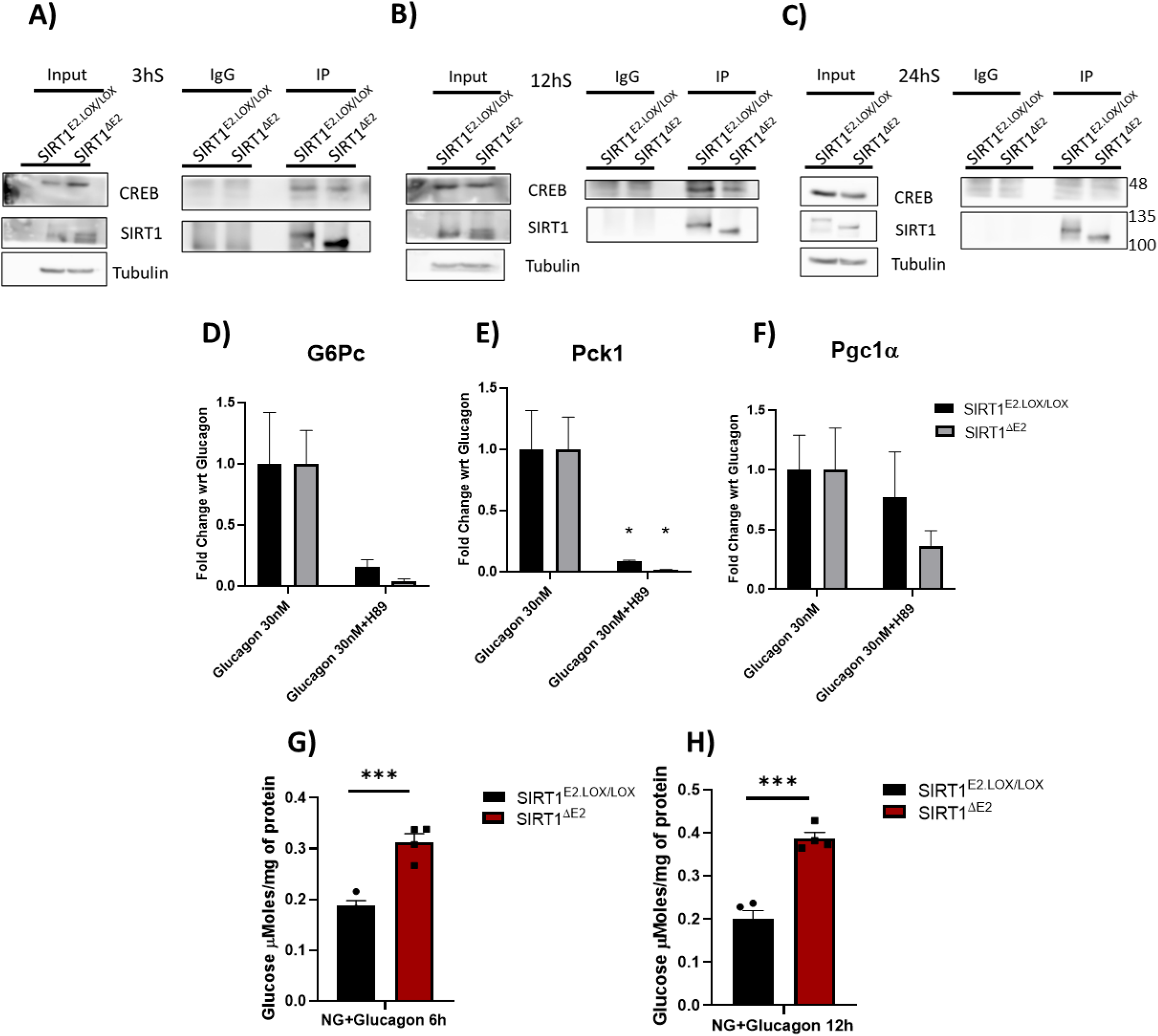
(Sup. Fig. 3): Dynamic Interaction of SIRT1 with CREB and Functional Consequences in Gluconeogenesis. (**A)-(C)** Time-resolved Co-IP with SIRT1 antibody from liver lysates collected after 3, 12, and 24 hours of starvation to assess dynamic SIRT1–CREB interaction in vivo. **(D)** Relative expression of G6pc, Pck1, and Pgc1α after treatment with a H89, PKA inhibitor. **(E)-(F)** Gluconeogenesis assay was performed in SIRT1^E2LOX/LOX^ and SIRT1^ΔE2^ hepatocytes cultured under various starvation conditions: NG + glucagon (6 h), and NG + glucagon (12 h). All graphs were plotted on GraphPad Prism 8.0, Student’s t-test was done for statistical analysis, significance as *, p<0.05; **, p<0.005; ***, p<0.0005.

**Supplementary Figure 4.**
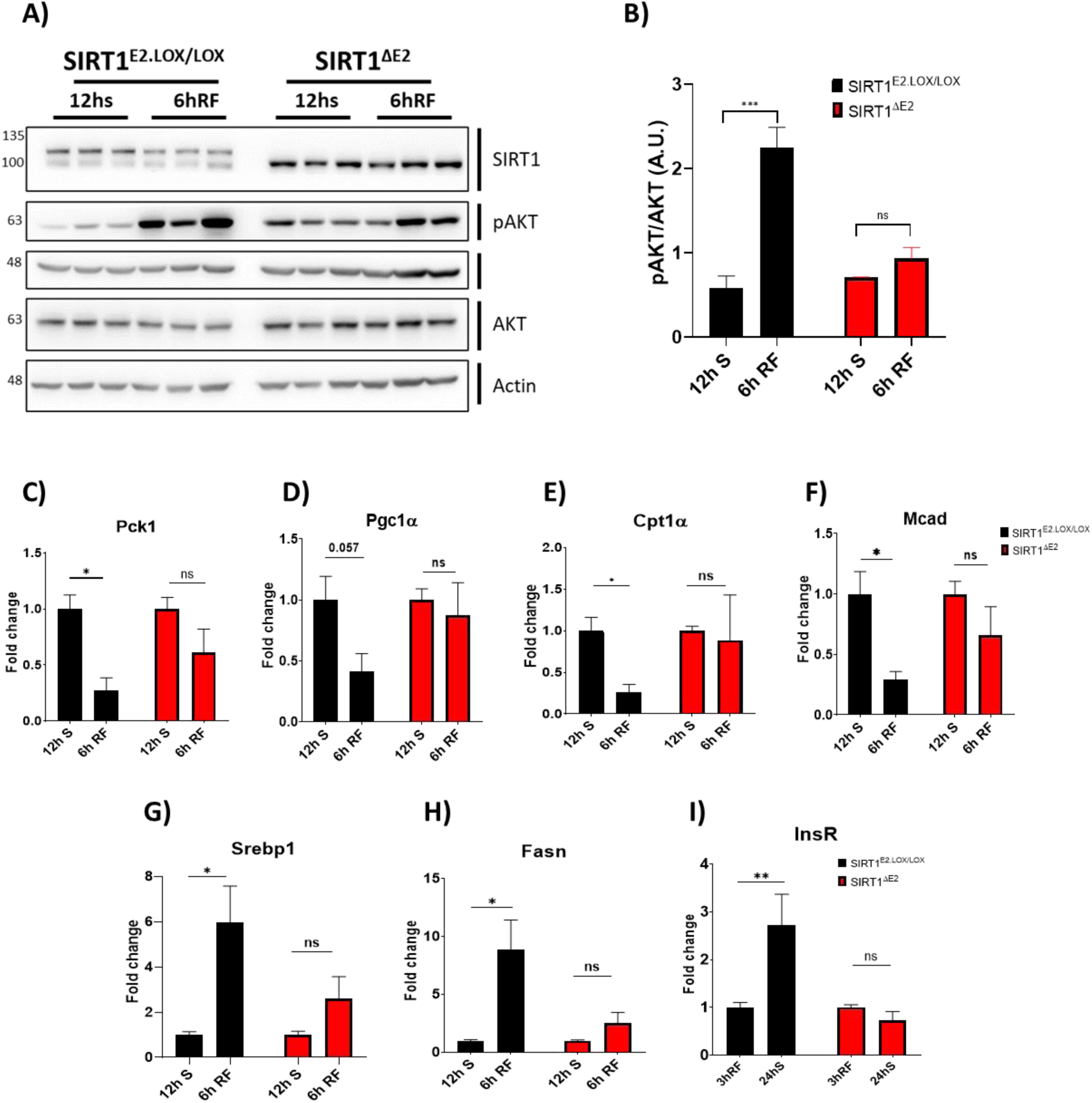
(Sup. Fig. 4): Altered Refeeding Response and Hepatic Gene Expression in SIRT1^ΔE2^ Mice. **(A)** Western blot analysis of liver lysates from 12-hour fasted and 6-hour refed SIRT1^ΔE2^ and SIRT1^E2.LOX/LOX^ male mice. Blots were probed for SIRT1, total AKT and phosphorylated AKT (Ser473). **(B)** Quantification of pAKT/AKT ratio normalised to loading control. **(C)-(I)** Relative mRNA expression of indicated hepatic genes from the same fasting-refeeding paradigm, except for InsR, in that the mice were fed refed for 3hrs. All graphs were plotted in GraphPad Prism 8.0, Student’s t-test and 2-way Anova were used for statistical significance, *, p<0.05; **, p<0.005; ***, p<0.0005.

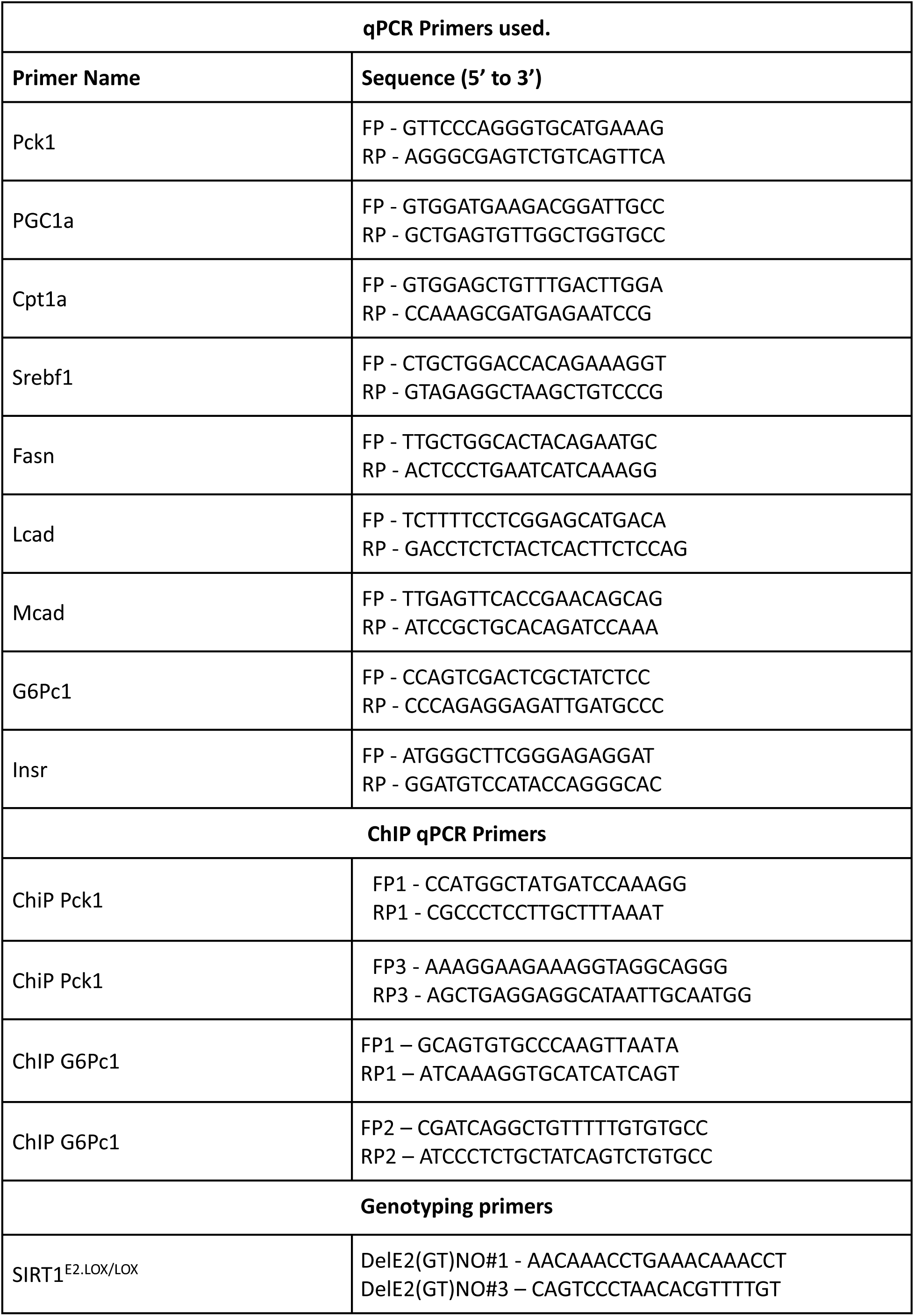
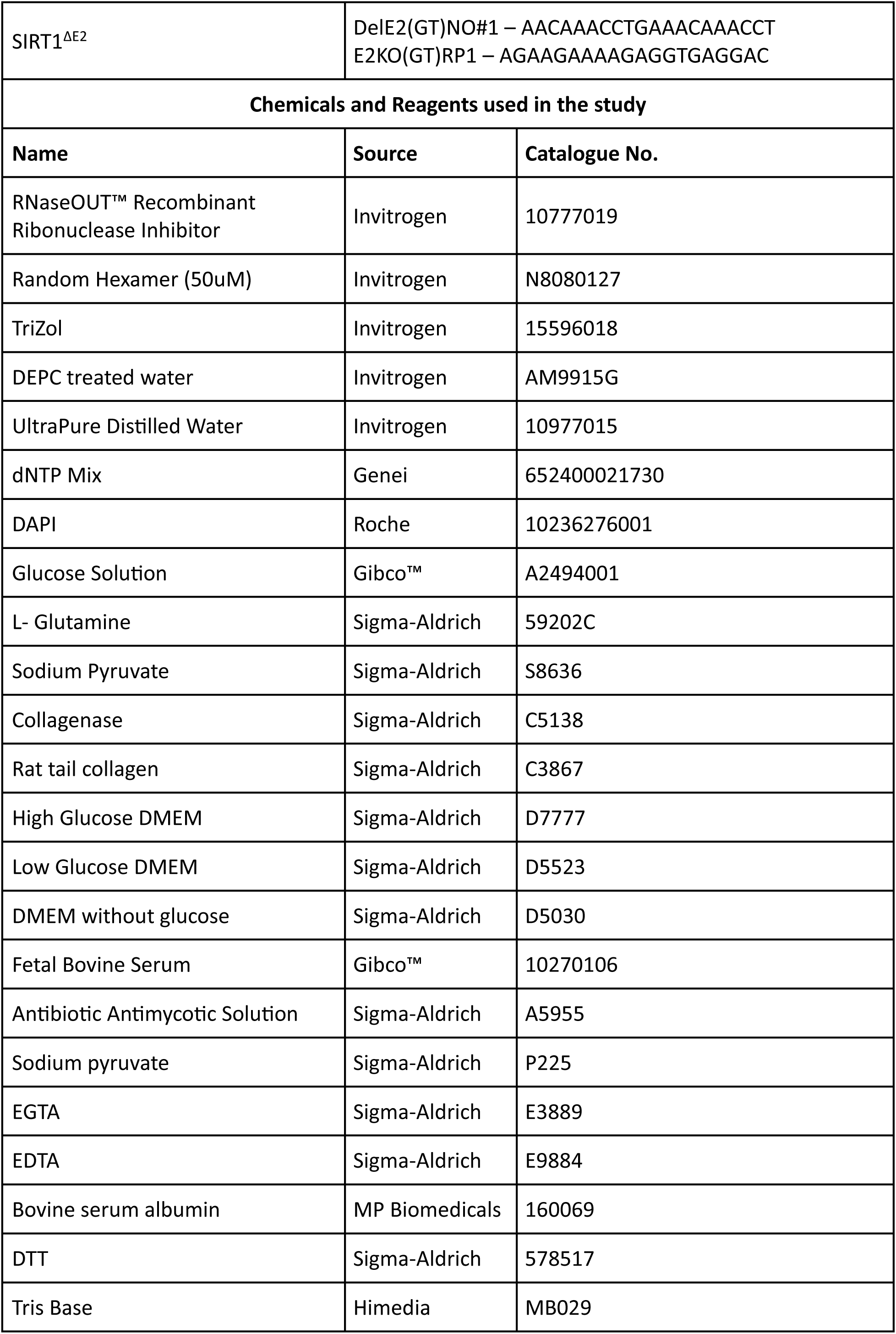
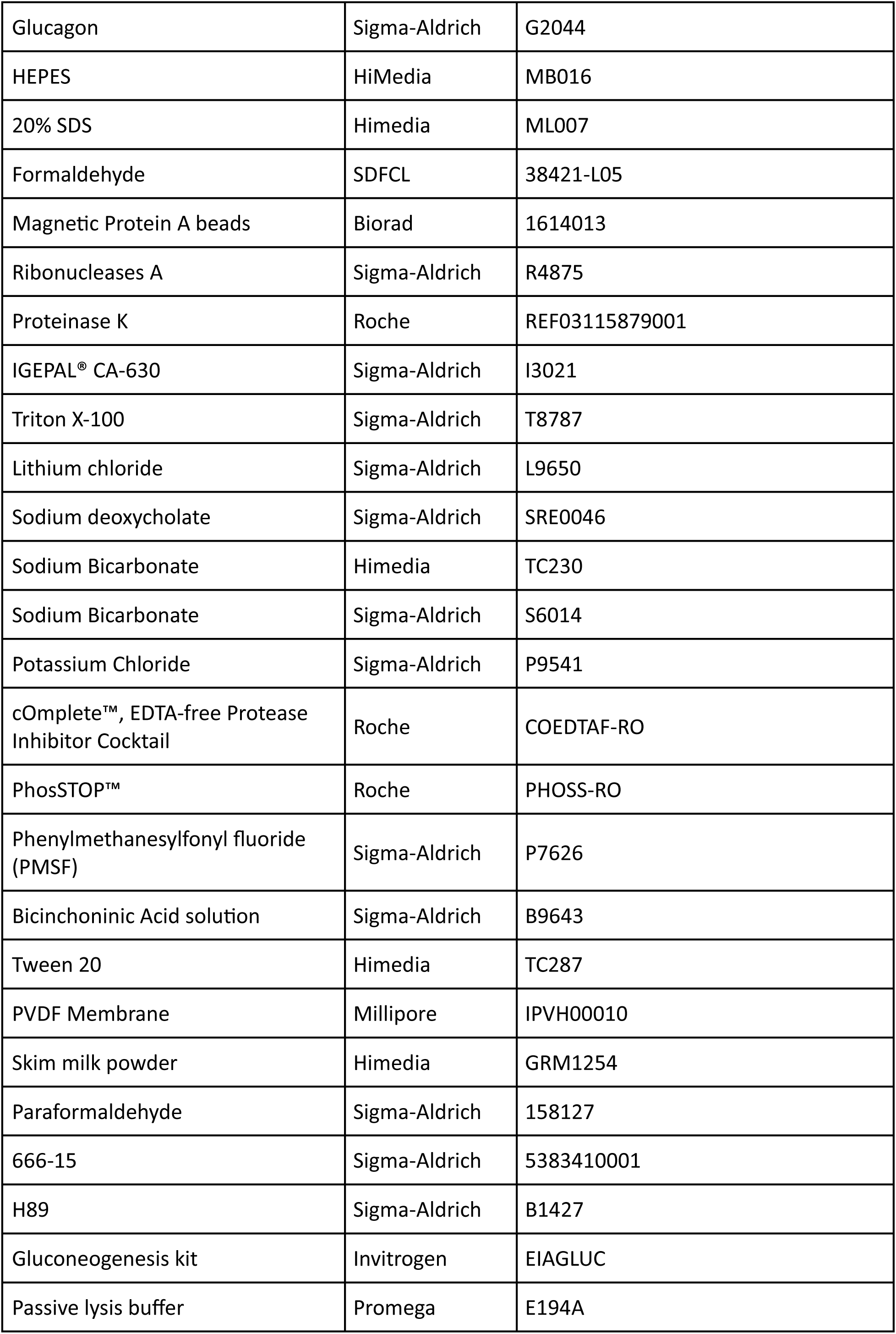
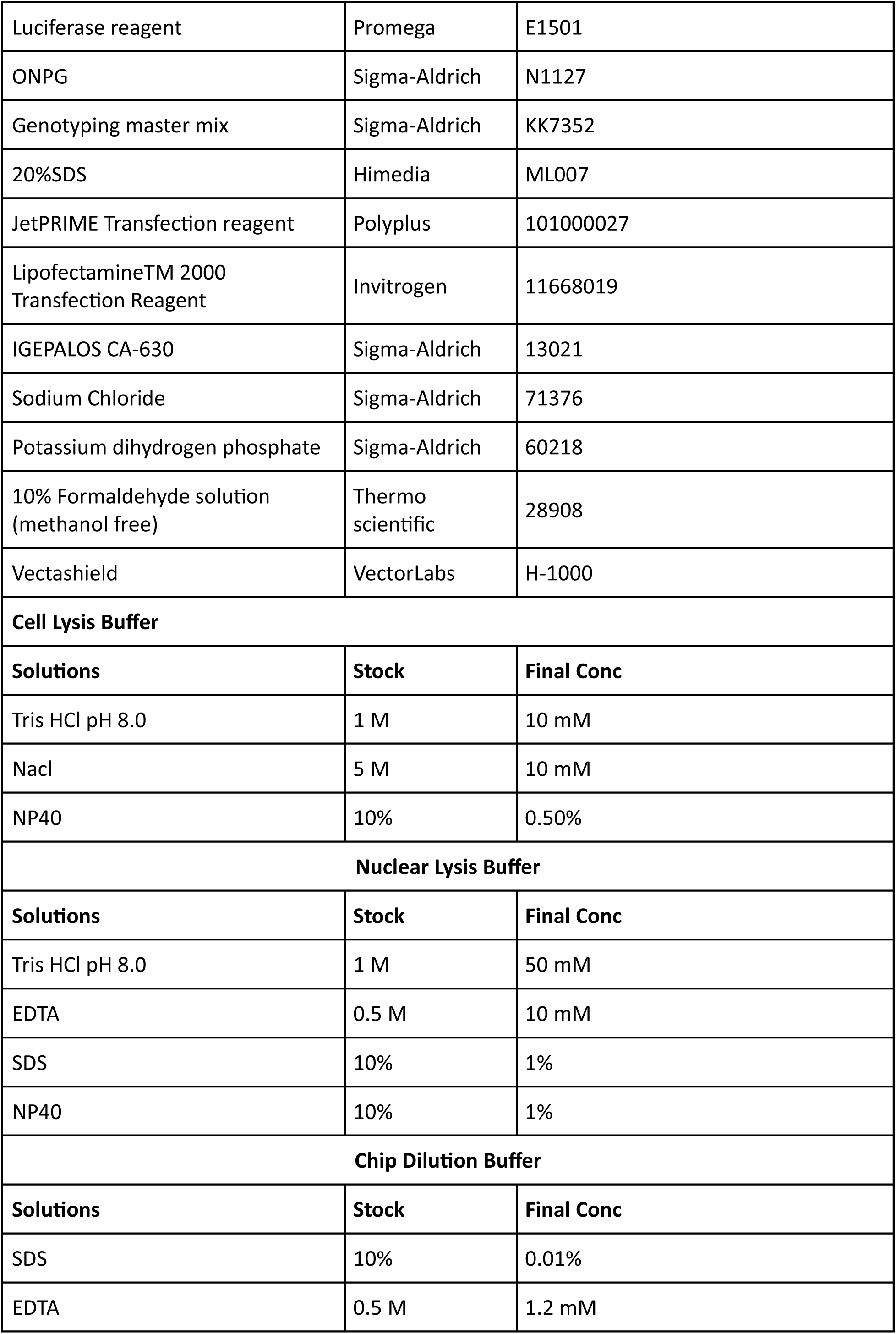
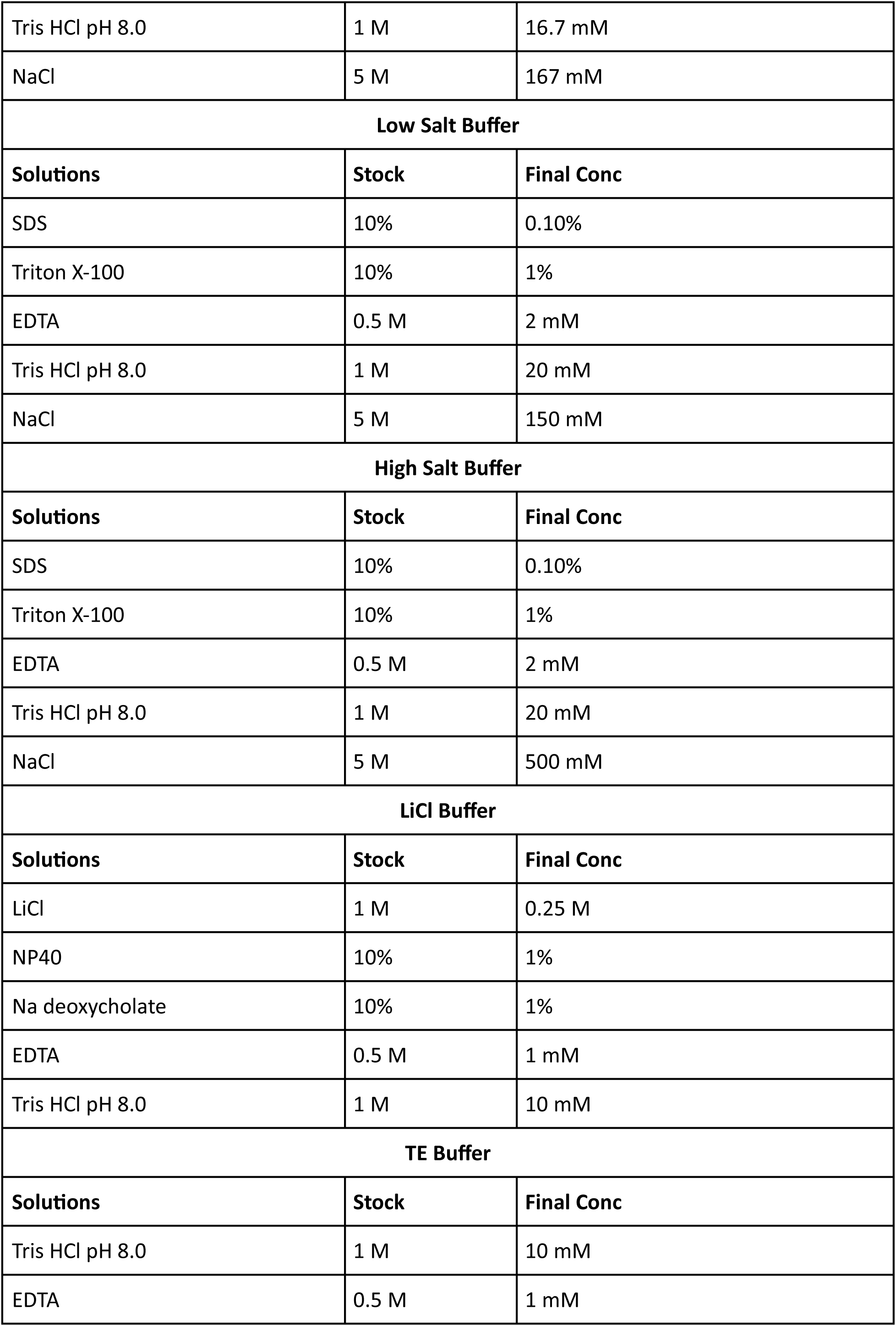
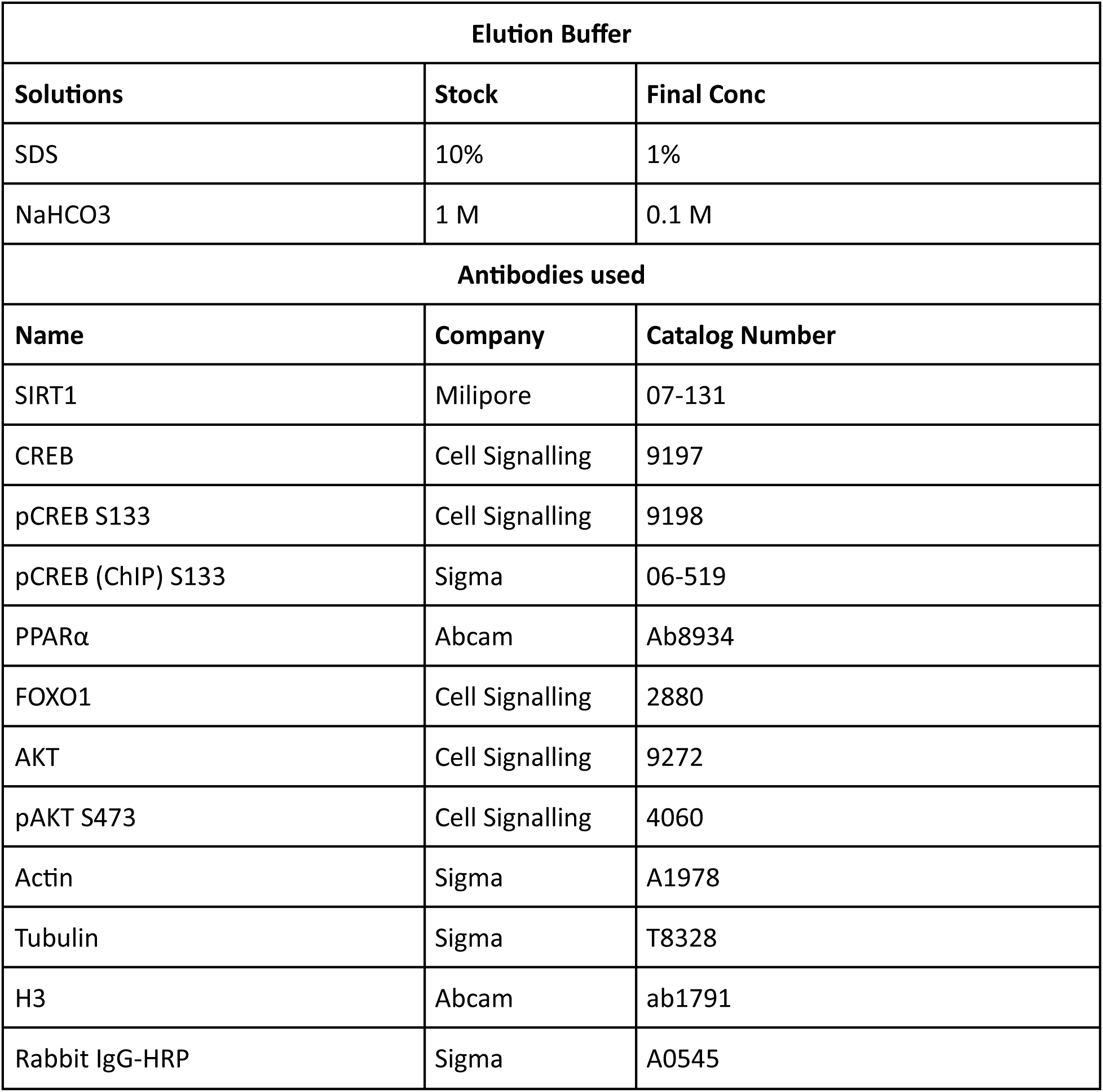

